# KRAS withdrawal in Cholangiocarcinoma leads to immune infiltration and tumor regression

**DOI:** 10.1101/2025.06.11.659210

**Authors:** Youwei Qiao, Matthew F. Yee, Chaitanya N. Parikh, Yueying Cao, Yu-Huan Shih, Boyang Ma, Joae Qiong Wu, Marcus Ruscetti, Shun-Qing Liang, Wen Xue

## Abstract

**Background and Aims:** Cholangiocarcinoma (CCA) is a liver cancer with poor survival rates. Current treatments, including targeted therapies for specific mutations, are limited and benefit only a small subset of patients. *KRAS* mutations are found in 15-40% of CCA, representing a new potential treatment target. Whether KRAS inhibition leads to CCA tumor regression is unknown partly due to the lack of conditional animal models.

**Approach and Results:** We engineered a conditional *Kras*^G12D^-driven CCA mouse model by co-delivering plasmids encoding the *Sleeping Beauty* transposase with a luciferase reporter, a transposon-borne inducible *Kras*^G12D^ transgene and Cas9 and *Trp53* guide RNA into mouse liver by hydrodynamic tail-vein injection. *In vivo* bioluminescent imaging showed that *Kras*^G12D^ withdrawal resulted in 99% tumor regression by day 7. *Kras*^G12D^ withdrawal resulted in infiltration of activated CD8^+^ T cells by IHC and IF staining. Single cell RNA-Seq result also validated the enrichment of activated CD8^+^ T cells subpopulation in *Kras*^G12D^-withdrawn tumor. RNA-Seq suggested that *Kras*^G12D^ withdrawal stimulated transforming growth factor beta pathway and induced senescence. We used cytokine array to characterize the secretion of pro-inflammatory factors, including IL-15 and Ccl17, upon *Kras*^G12D^ withdrawal. Lentiviral overexpression of murine CCL17 delayed CCA tumor progression in a xenograft model, and overexpression of murine IL-15 resulted in tumor regression in a transplant model. Flow cytometry analysis revealed that IL-15 and CCL17 recruited and activated CD8^+^ T cells in CCA tumor. Expression of IL-15 resulted in blockade of tumor progression in our TKP CCA model.

**Conclusions:** *Kras*^G12D^ withdrawal results in rapid tumor regression, highlighting the importance of oncogenic *Kras* in CCA tumor maintenance. *Kras*^G12D^ withdrawal induces p53-independent senescence, secretion of pro-inflammatory factors, and immune surveillance by activated CD8^+^ T cells. We identified two secreted factors IL-15 and CCL17 that could recruit and activate T cells and control CCA tumor progression. This study underscores KRAS inhibition as a potential therapeutic approach for CCA.

## Introduction

Cholangiocarcinoma (CCA) is the second most prevalent type of primary liver cancer, which accounts for roughly 15% liver tumors (1–3). Currently, surgical resection is the preferred treatment for CCA (1–3), but most patients are diagnosed at an advanced stage with unresectable disease, due to a lack of tumor markers for early diagnosis. The prognosis of CCA is therefore dismal, with a 5-year survival rate of 10% (1–3). Targeted molecular therapies for CCA are in clinical trials, including isocitrate dehydrogenase 1 (IDH1) inhibitors and fibroblast growth factor receptor 2 (FGFR2) inhibitors (1, 4, 5). But these targeted molecular therapies can only benefit a small percentage of patients due to the low prevalence of IDH1 and FGFR2 mutations in CCA. Thus, there is an urgent need to develop novel targeted molecular therapies that benefit more CCA patients.

*KRAS* is one of the most frequently mutated proto-oncogenes in cholangiocarcinoma. Fifteen to forty percent of patients with CCA harbor a *KRAS* mutation (1–3, 6–8). KRAS was regarded as an “undruggable” target for decades due to the lack of binding pockets for small molecule inhibitors on its “smooth” surface (7, 9). Recently, selective, potent inhibitors for mutant KRAS have been developed (10–12). In 2021, the U.S. Food and Drug Administration granted accelerated approval for KRAS(G12C) inhibitor “sotorasib” for non-small cell lung cancer treatment (13). Therefore, targeting *KRAS* mutations has become a potential therapeutic approach for CCA patients with advanced disease. However, there are limited *in vivo* models to study the mechanism by which *KRAS* mutations promote tumor maintenance in CCA. Thus, it is crucial to conduct animal studies to explore the therapeutic potential of KRAS inhibition for cholangiocarcinoma treatment.

*KRAS* is a member of rat sarcoma viral oncogenes (RAS) family, which encodes a GTPase that functions as a molecular switch regulating intracellular signaling cascades by switching between activated and inactivate conformation (10, 11, 14). Oncogenic mutations result in a constitutively activated KRAS and contribute to cancer cell proliferation, tumor metabolism, and resistance to cell death and cell cycle arrest (15–17). A previous study has shown that oncogenic *Kras* withdrawal induces apoptotic cell death in an inducible *Kras*-driven lung adenocarcinoma mouse model and pancreatic ductal adenocarcinoma (PDAC) models (18–20). However, many questions remain poorly understood, including whether KRAS inhibition leads to CCA tumor regression, what are the mechanisms of tumor regression and what role immune system plays in this process.

In this study, we investigated the role of oncogenic *Kras* in tumor maintenance in an inducible *Kras*-driven CCA mouse model. We developed a conditional CCA mouse model by codelivery of plasmids encoding a transposon-borne inducible *Kras*^G12D^ transgene and Cas9 and *p53* guide RNA. We show that *Kras*-driven CCA tumors regress upon withdrawal of oncogenic *Kras*, accompanied by activation of transforming growth factor beta pathway and cellular senescence. Senescent CCA tumor cells secreted pro-inflammatory factors including IL-15 and CCL17, and lentiviral overexpression of either *Il-15* or *Ccl17* delayed Ras-driven CCA xenograft tumor progression. Consistent with results in the transplant model, expression of *Il-15* delayed tumor progression in our TKP CCA model.

## Results

### Conditional overexpression of Kras^G12D^ with Trp53 knockout drives the formation of Cholangiocarcinoma in B6 mice

A cancer genomic study of 195 patients suggests that *KRAS* and *TP53* are among the most frequently altered genes in cholangiocarcinoma (CCA) (7) (**Fig. S1A**). According to this study, thirty one percent of patients harbor *TP53* mutations, and sixteen percent of patients harbor *KRAS* mutations (**Fig. S1B**). Among CCA patients which harbor *KRAS* mutations, over forty percent carry *KRAS*^G12D^ mutations (21) (**Fig. S1C**). In mice, liver-specific expression of *Kras*^G12D^ with deletion of *Trp53* gene drives the formation of liver tumors that resemble CCA both histologically and molecularly—e.g., activation of MAPK/ERK and PI3K/AKT pathways (22–24). To engineer an inducible model of CCA, we delivered a *Sleeping Beauty* transposon plasmid encoding Tet-On-inducible *Kras*^G12D^ transgene and reverse-tetracycline transactivator (rtTA), a plasmid encoding Cas9 nuclease and *Trp53* guide RNA (25–27), and a plasmid encoding *Sleeping Beauty* transposase and luciferase reporter to mouse liver by hydrodynamic tail-vein injection (**Fig. 1A**). Oncogenic *Kras*^G12D^ will be overexpressed in livers of mice fed a diet containing doxycycline (*Kras* ON) and silenced in livers of mice fed on a normal diet (*Kras* OFF) (**Fig. 1A**). We named this mouse model TRE.*Kras*^G12D^/*p53* knockout CCA (TKP).

**Fig. 1.**
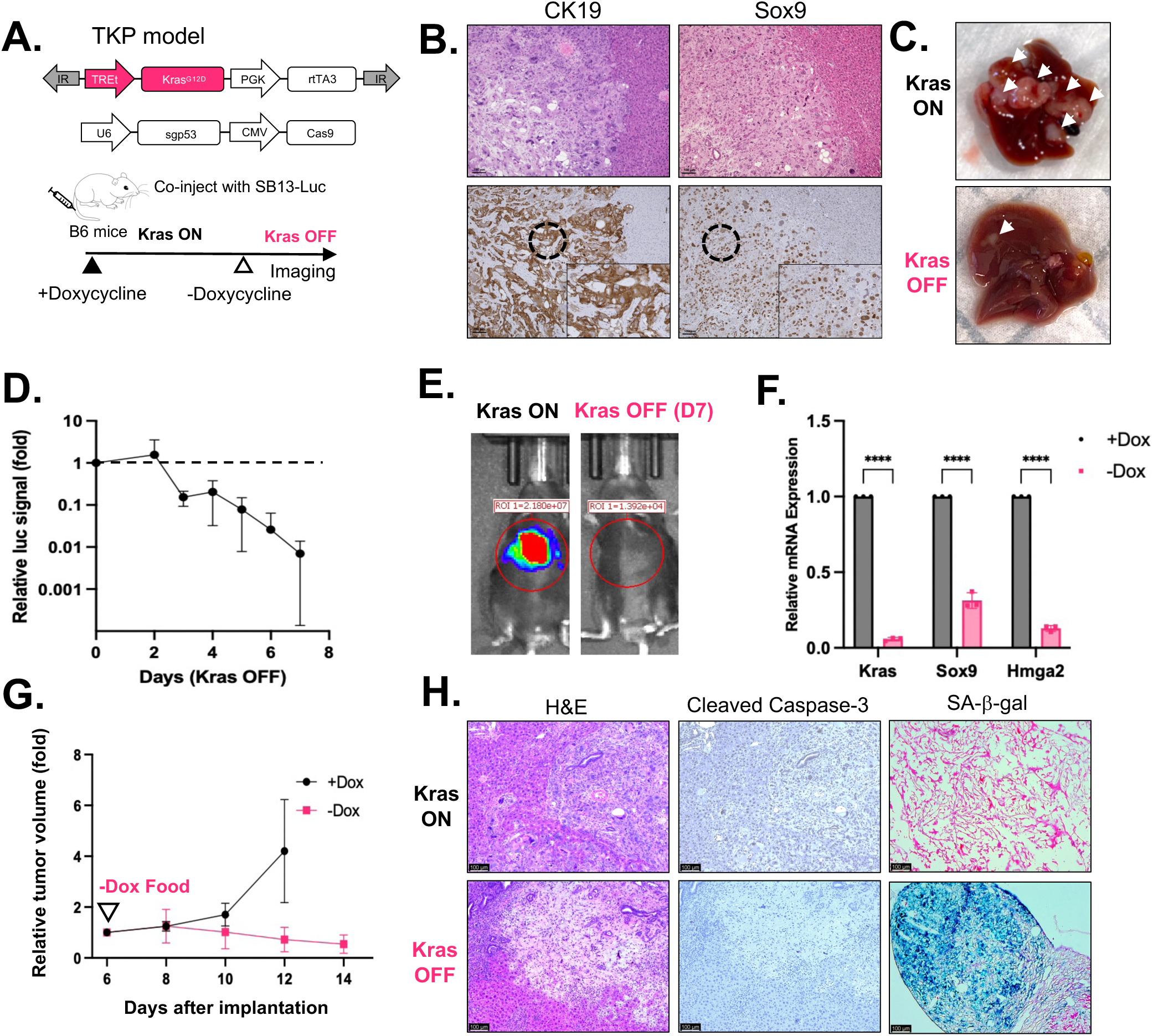
Oncogenic *Kras*^G12D^ withdrawal in CCA mouse model induces tumor regression and senescence. (**A**) Generating a doxycycline-inducible CCA mouse model using *sleeping beauty* transposon and CRISPR/Cas9 (TRE.*Kras*^G12D^;*Trp53* KO, TKP model). Plasmids encoding transposase with luciferase reporter, TRE.Kras^G12D^ transposon, and Cas9 with sgRNA targeting *Trp53* were hydrodynamically injected into immunocompetent B6 mice. **(B)** Hematoxylin & eosin (H&E) and immunohistochemistry (IHC) staining of Kras ON tumors 4 weeks after injection. CK19, Cytokeratin 19. Sox9, SRY-Box Transcription factor 9. Scale bar=100µm. **(C)** Representative gross liver images of TKP *Kras* ON and TKP *Kras* OFF mice. Arrows indicate tumors. **(D)** Relative luciferase signal of liver tumors demonstrates tumor regression. Day 0 (*Kras* ON) is set as 1. Data are presented as mean ± SD. (n=3). **(E)** Representative luciferase imaging shows *Kras* OFF tumor regression. **(F)** qRT-PCR analysis of *Kras*, *Sox9*, *Hmga2* mRNA expression in TKP cell line on Dox or off Dox at day 8. Each group has three biological replicates. Data are presented as mean ± SD. P-values were calculated by unpaired t-test. \*\*\*\**P* <0.0001. **(G)** Change in tumor volume of mice bearing subcutaneous TKP tumors. Tumor volume at day 6 is set as 1. Data are presented as mean ± SD. (n=4 for +Dox group and n=6 for –Dox group). **(H)** H&E staining and IHC staining and senescence-associated beta-galactosidase (SA-β-Gal) staining of TKP *Kras* ON and TKP *Kras* OFF tumors. *Kras* OFF tumors are positive for SA-β-Gal and negative for Cleaved Caspase-3. Scale bars are 100µm.

Consistent with a previous study using constitutive expression of *Kras*^G12D^ (24), inducible overexpression of *Kras*^G12D^ with p53 knockout drove the formation of liver tumors that were CK19-positive, a key diagnostic marker of CCA (28) (**Fig. 1B**). The tumor additionally showed elevated SOX9 expression by IHC, a transcription factor linked to poor prognosis in human CCA, highlighting the aggressiveness of our tumor model (29) (**Fig. 1B**). The earliest visible surface tumors appeared around 2 weeks post-injection; luciferase signal could be detected at 3 weeks; and at 4 weeks post-injection, *Kras* ON mice developed extensive liver tumors and were hunched and sick (**Fig. 1C, *Kras* ON**). The majority of tumor-bearing mice succumbed within six weeks post-injection. These findings suggest that our tumor mouse model resembles aggressive human CCA tumors with poor prognosis.

### Oncogenic Kras^G12D^ withdrawal results in cellular senescence and tumor regression in vivo

The TKP model allows us to study the effects of *Kras*^G12D^ withdrawal. To confirm the conditional expression of *Kras*, we isolated TKP cell line from a *Kras* ON solid tumor 4 weeks post-injection and maintained the TKP cell line in medium with doxycycline (+Dox). Consistent with previous studies (30), *Kras* ON cells expressed high levels of liver progenitor cell markers Hmga2 and Sox9 (**Fig. 1F**). High expression level of HMGA2 is associated with poor survival in liver cancer patients (30). Oncogenic *Kras* withdrawal by removing doxycycline (-Dox) resulted in a dramatic downregulation of *Kras*, *Sox9*, and *Hmga2* expression in the TKP cell line (**Fig. 1F**).

To investigate whether *Kras*^G12D^ withdrawal leads to tumor regression in the inducible CCA mouse model, we removed doxycycline diet at ∼4 weeks post-injection (**Fig. 1A**). Bioluminescent imaging of mice revealed a steady decline in tumor size to an ∼99% reduction by day 7 (**Fig. 1D,E** and **Fig. S2**). Surface liver tumors were significantly reduced at 8 days after doxycycline withdrawal (**Fig. 1C, *Kras* OFF**). To test whether tumors regress upon *Kras* withdrawal in a transplant model, we injected the TKP cell line subcutaneously into B6 mice fed on doxycycline diet, and after tumor onset (6 days post-injection), we removed doxycycline from the diet. Whereas xenograft tumors continued to grow in mice that remained on doxycycline, xenograft tumors shrank in mice taken off doxycycline, validating that oncogenic *Kras* withdrawal results in tumor regression (**Fig. 1G**).

Oncogenic *KRAS* plays a vital role in regulating cell death and cell cycle arrest mechanisms, promoting cancer cell proliferation (15–17). To determine if tumor regression results from cell death or cell cycle arrest, we performed IHC staining on *Kras* ON tumors and day 8 *Kras* OFF tumors. *Kras* OFF tumors were negative for Cleaved-Caspase 3, indicating that apoptotic cell death is not the mechanism of tumor regression upon Kras withdrawal (**Fig. 1H**). *Kras* OFF tumors were positive for senescence-associated beta-galactosidase (SA-β-gal) activity (**Fig. 1H**), suggesting that oncogenic *Kras* withdrawal results in cellular senescence.

Consistent with the senescent phenotype of *Kras* OFF tumors, *Kras* OFF TKP cells showed abrogated cell growth (**Fig. 2A**, *Kras* OFF –Dox), even upon reactivation of *Kras*^G12D^ expression by doxycycline treatment (**Fig. 2A**, *Kras* OFF +Dox), compared with *Kras* ON TKP cells (**Fig. 2A**, *Kras* ON +Dox). *Kras* OFF TKP cells were positive for SA-β-gal activity and exhibited a significant altered morphology compared to proliferating *Kras* ON cells, characterized by an enlarged, flattened appearances with expanded nuclei (**Fig. 2B**). Additional molecular markers of cellular senescence—p15, p16, and p27—were upregulated in *Kras* OFF TKP cells compared to *Kras* ON TKP cells (**Fig. 2C**), indicating that oncogenic *Kras* withdrawal results in cell cycle arrest and cellular senescence.

**Fig. 2.**
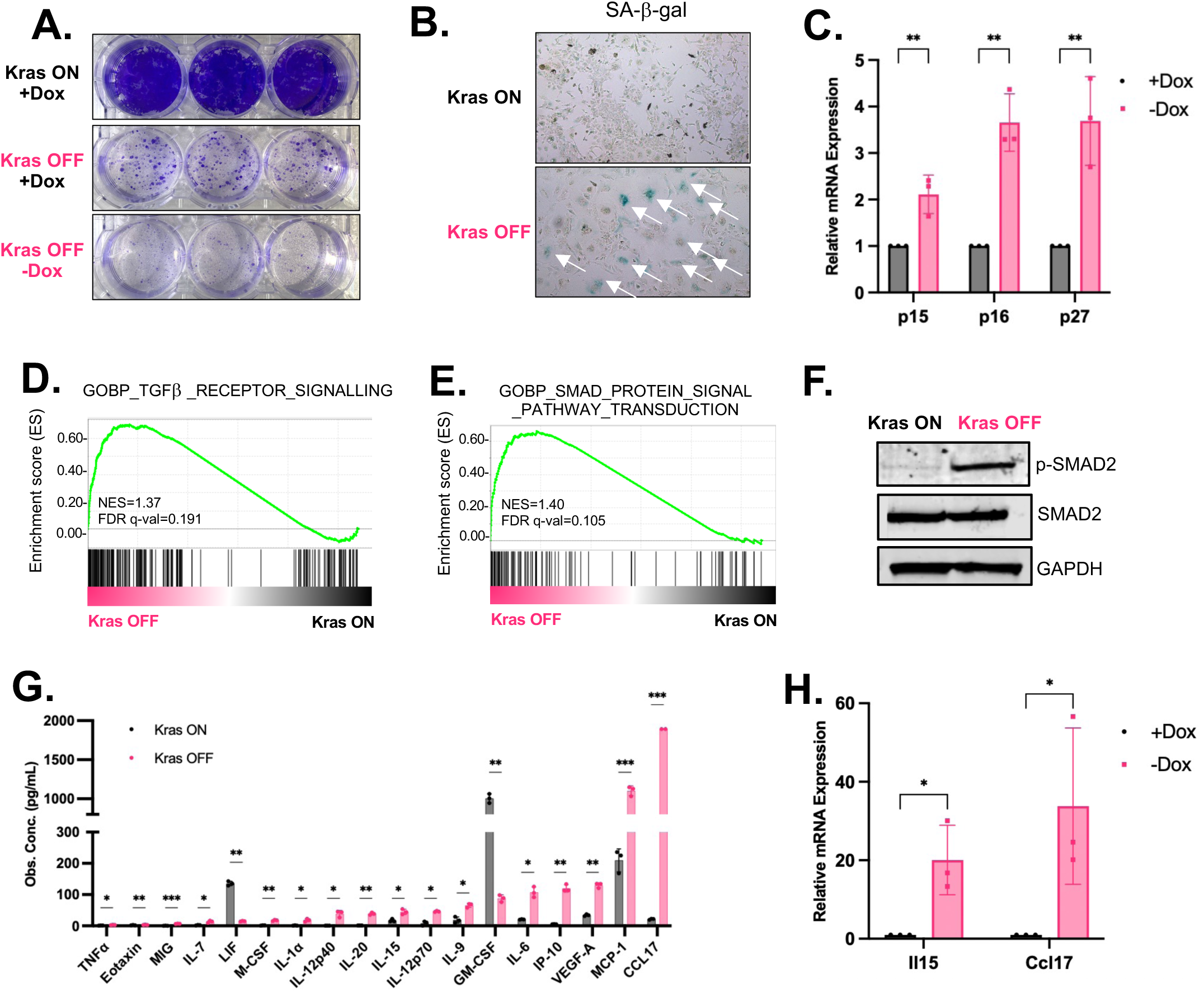
*Kras*^G12D^ withdrawal results in cellular senescence and alters the secretory phenotype. **(A)** Colony formation assay of TKP *Kras* ON and *Kras* OFF cell. *Kras* OFF (+Dox) TKP cells indicated *Kras* OFF TKP cells which were re-introduced with doxycycline. Each group has three technical replicates. **(B)** SA-β-Gal staining of TKP *Kras* ON and *Kras* OFF cells. Arrows indicate senescent cells. **(C)** qRT-PCR analysis of p15, and p16, p27 mRNA expression in TKP cell line on Dox or off Dox at day 8. Each group has three biological replicates. **(D-E)** GSEA analysis in TKP *Kras* OFF and *Kras* ON cell lines. NES, normalized enrichment score. FDR, false discovery rate. **(F)** Phospho-SMAD2, SMAD2, and GAPDH western blot in TKP *Kras* ON and *Kras* OFF cell line. Cells were starved with 0.2% FBS. **(G)** Cytokine array of TKP *Kras* ON and *Kras* OFF cell line. Each group have three biological replicates. One outlier from Ccl17 group is omitted. **(H)** qRT-PCR analysis of Il15 and Ccl17 in Kras ON and Kras OFF TKP cell line. Each group has three biological replicates. (B, H) Data are presented as mean ± SD. P-values were calculated by unpaired t-test. (G) Data are presented as mean ± SD. P-values were calculated by paired t-test. \**P* <0.05, \*\**P* <0.01, \*\*\**P* <0.001, \*\*\*\**P* <0.0001.

### Kras^G12D^ withdrawal changes the transcriptional landscape of TKP CCA cell line

RNA-Seq analyses revealed that *Kras*^G12D^ withdrawal significantly altered the transcriptional landscape of TKP CCA cells (**Fig. S3A**). Pearson correlation analysis between RNA-Seq replicates revealed strong positive correlations, supporting the reliability of the datasets for differential expression analyses (**Fig. S3B**). As expected, *Kras* transcript levels (reads-per-kilobase per million mapped reads; RPKM) were significantly reduced by approximately 5-fold in *Kras* OFF TKP cells (**Fig. S3C**). Among all differentially expressed genes, we identified 1,430 genes significantly upregulated and 644 genes significantly downregulated (≥2-fold; **Fig. S3D**). Interestingly, genes downregulated upon *Kras*^G12D^ withdrawal included *Lilr4b* and *Cxcr2* (31–33) (**Fig. S3A**). Blockade of CXCR2 has been shown to promote anti-tumor immunity, suggesting that *Kras*^G12D^ withdrawal might trigger immune response (34, 35). Gene set enrichment analysis (GSEA) suggests that pathways related to transforming growth factor beta (TGF-β) signaling (**Fig. 2D and Fig. S4A**) and SMAD protein signaling (**Fig. 2E and Fig. S4B-D**) were among the top enriched gene sets upon *Kras*^G12D^ withdrawal in TKP cells. Indeed, *Tgfb2* levels were markedly upregulated in the RNA-seq data (**Fig. S3A**). Consistent with the upregulation of TGF-β–SMAD signaling, we detected increased levels of phosphorylated SMAD2 (p-SMAD2) in *Kras* OFF TKP cells compared to *Kras* ON TKP cells, whereas total SMAD2 protein levels were the same (**Fig. 2F**). A previous study has shown that treating liver cancer cells with TGF-β activates SMAD protein signaling pathway, upregulates cell cycle inhibitors p15 and p21, and induces cellular senescence (36). Similarly, our results show that oncogenic *Kras* withdrawal inhibits TKP cell proliferation, activating TGF-β and SMAD signaling pathway, and inducing cellular senescence.

### Oncogenic Kras^G12D^ withdrawal alters the secretory phenotype of TKP CCA cell line

Cellular senescence arrests tumor growth by downregulating proliferation genes and upregulating genes encoding anti-tumorigenic secretory factors known as senescence-associated secretory phenotype (SASP) (37–40). SASP factors often include proinflammatory chemokines and cytokines that recruit and activate tumor-fighting immune cells, angiogenic factors, growth factors, and matrix metalloproteinases (37–40). RNA-Seq analysis revealed that *Kras*^G12D^ withdrawal altered the transcription level of essential cell cycle and senescence-associated secretory phenotype (SASP) genes, indicating that oncogenic *Kras* withdrawal induces cell cycle arrest and a pro-inflammatory, senescent phenotype (**Fig. S3E-F**). Cytokine array analyses of *Kras* OFF TKP cells and *Kras* ON TKP cells revealed that *Kras*^G12D^ withdrawal stimulates the production of proinflammatory chemokines involved in NK cell recruitment (IP-10/CXCL10 and MCP-1/CCL2), cytokines involved in NK cell and CD8^+^ T cell activation and proliferation (IL-12, IL-15, TNF-α), and angiogenic factors (VEGF) (38, 39) (**Fig. 2G and Fig. S5A**). A previous study has shown that IL-15 can suppress metastatic and primary liver cancer by activation of CD8^+^ T cells (41). Interestingly, CCL17, a chemokine that has not been extensively studied in liver cancer, increased dramatically upon *Kras*^G12D^ withdrawal in TKP cells (**Fig. 2G and Fig. S5A**). A previous work has shown that CCL17 modulates local immune environment in lung adenocarcinoma (42). *Il15* and *Ccl17* were significantly upregulated upon *Kras*^G12D^ withdrawal in our RNA-Seq analyses and by RT-qPCR (**Fig. 2H and Fig. S3A**). These results indicate that *Kras*^G12D^ withdrawal in TKP cells induces SASP factors, including a well-studied cytokine in liver cancer (IL-15) and previously uncharacterized chemokine (CCL17).

Previous studies have shown that SASP involves activation of the NF-kB pathway, which can induce expression of IL-15 and CCL17 (43). Our GSEA results further indicate enrichment of pathways associated with inflammatory and immune signaling upon *Kras*^G12D^ withdrawal, including interferon alpha and gamma pathways (**Fig. S5B, C**). To confirm these observations, we performed qRT-PCR analysis and observed increased mRNA levels of IL-15 when we treated TKP *Kras* ON cells with interferon alpha and gamma (**Fig. S5D, E**). These results support our hypothesis that IL-15 is regulated by interferon signaling pathways.

### Kras^G12D^ withdrawal alters cellular subpopulations and recruits CD8 T cells to tumors

We next explored whether SASP induction triggered by oncogenic *Kras* withdrawal activates CD8^+^ T cells. In histological analyses of *Kras* ON tumors and day 8 *Kras* OFF tumors, we detected a dramatic increase in CD8^+^ T cells and F4/80^+^ macrophages in *Kras* OFF tumors compared to *Kras* ON tumors (**Fig. 3A, B**), suggesting that *Kras*^G12D^ withdrawal from CCA tumors promotes an immune response. Immunofluorescence analysis of CK19, CD8, and activation marker CD69 revealed notable differences in T cell distribution and activation state in *Kras* ON and *Kras* OFF TKP tumors. In *Kras* ON tumors, CD8 T cells were sparsely distributed in CK19-positive CCA tumor cells (**Fig. 3C, *Kras* ON**). However, in *Kras* OFF tumors, there was a marked increase in density of CD8 and CD69 double positive T cells clustering around CK19-positive tumor cells (**Fig. 3C, *Kras* OFF**).

**Fig. 3.**
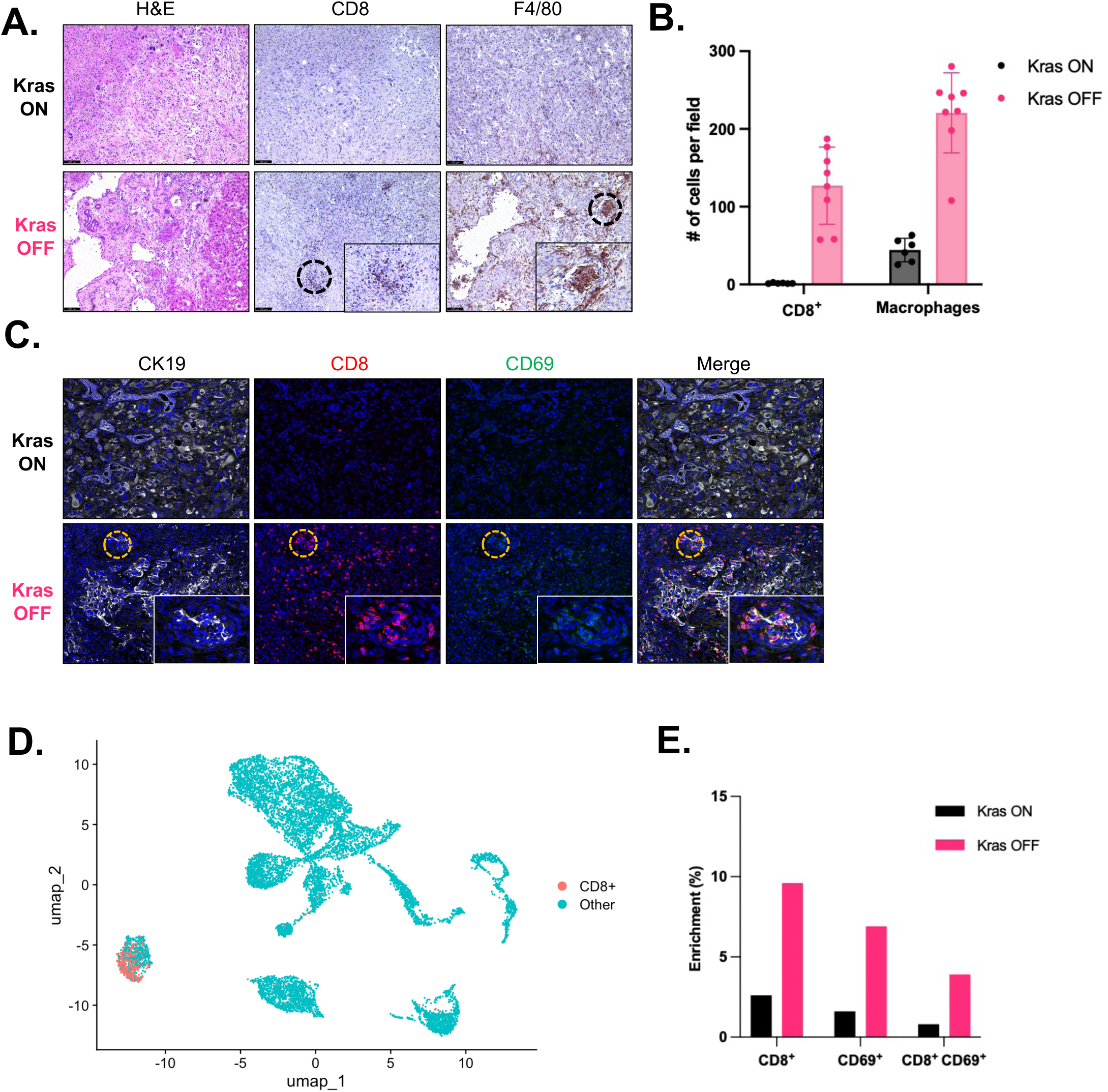
*Kras*^G12D^ withdrawal recruits CD8 T cells and Macrophages to tumor. **(A)** H&E and IHC staining of TKP *Kras* ON and *Kras* OFF tumors. *Kras* OFF tumors were collected 8 days after doxycycline withdrawal. **(B)** Quantification of CD8 and F4/80 in TKP *Kras* ON tumors (n=6) and *Kras* OFF tumors (n=8). Each data points represents a mean of positive cells of ten 20X microscopic field. Lines represent mean value. P-values were calculated by unpaired t-test. \*\*\**P* <0.001. **(C)** Representative immunofluorescence images of CK19, CD8, and CD69 positive cells in TKP *Kras* ON and *Kras* OFF tumors. **(D)** UMAP visualization of CD8^+^ T cell clusters identified by scRNA-seq of 13,329 cells from one TKP *Kras* ON and one TKP *Kras* OFF tumors. **(E)** Bar graph indicating the enrichment of CD8^+^ T cells, CD69^+^ T cells, CD8^+^ CD69^+^ T cells in TKP *Kras* ON and *Kras* OFF tumors by scRNA-seq.

To investigate the cellular diversity within TKP tumors and the effects of *Kras*^G12D^ withdrawal, we performed single-cell RNA sequencing (scRNA-seq) on cells isolated from one TKP tumor with sustained *Kras*^G12D^ expression (*Kras* ON) and one TKP tumor following *Kras*^G12D^ withdrawal (*Kras* OFF). We obtained transcriptomic profiles from a total of 13,329 cells, which were assigned to 15 distinct subclusters using UMAP dimensionality reduction and Louvain clustering (**Fig. S6A, B**). Cell types were identified based on the expression of established marker genes, including Ptprc for immune cells, *Csf1r*, *C1qa*, *Cd68*, and *Ccr2* for macrophages, *S100a8* and *Cxcr2* for neutrophils, *Cd3d* and *Cd3e* for T cells, *Pecam1* and *Flt1* for vascular cells, *Xcr1* and *Cd209a* for dendritic cells, *Krt8*, *Krt18*, and *rtTA3* for tumor cells, *Dcn*, *Col1a1*, and *Acta2* for cancer-associated fibroblasts (CAFs), and *Cd79a*, *Cd19*, and *Igkc* for B cells (**Fig. S6C**).

*Kras*^G12D^ withdrawal significantly altered the cellular composition of the tumor microenvironment. Specifically, T cells, B cells, vascular cells, and CAFs were significantly enriched in *Kras* OFF tumors compared to *Kras* ON tumors (**Fig. S6D, E**). To further explore the T cell compartment, we re-clustered the T cell population and identified subpopulations of CD8^+^ T cells (expressing CD8a) and activated T cells (expressing CD69). Our analysis revealed that CD8^+^ T cells, CD69^+^ T cells, and double-positive CD8^+^ CD69^+^ T cells were significantly more abundant in *Kras* OFF tumors (**Fig. 3D, E**). This observation was corroborated by co-immunofluorescence (co-IF) staining, which similarly showed increased infiltration of CD8^+^ and CD69^+^ cells in *Kras* OFF tumors (**Fig. 3C**). These findings suggest that *Kras*^G12D^ withdrawal may enhance anti-tumor immunity by promoting the infiltration and activation of CD8^+^ T cells within the tumor microenvironment.

### Lentiviral expression of mouse Il-15 or Ccl17 delays Ras-driven CCA xenograft tumors

A previous study has shown that IL-15 suppresses tumor growth in a toxin-induced HCC mouse model, and the function of CCL17 in liver cancer remains unclear (41, 42). To characterize the functions of IL-15 and CCL17 *in vivo* in a CCA mouse model, we expressed mouse *Il-15* or *Ccl17* using lentiviral vector in a syngeneic Ras-driven CCA cell line, RIL-175, derived from liver tumors formed in C57BL/6 by intrasplenic injecting fetal p53^-/-^ hepatoblasts transduced with retroviral Hras^v12^ vectors (44). We transduced RIL-175 with lentiviral vectors encoding mouse *Il-15* or *Ccl17* cDNA or control empty lentiviral vector (**Fig. 4A**). RT-qPCR analyses and cytokine arrays confirmed that IL-15 and CCL17 were overexpressed and secreted in each cell line compared to control (**Fig. S7A-D**). Overexpression of IL-15 or CCL17 did not affect cell growth *in vitro* (**Fig. S7E**). We subcutaneously injected IL-15-expressing, CCL17-expressing, or control RIL-175 cells into immunocompetent C57BL/6 mice to initiate a syngeneic transplant mouse model of CCA. Compared to the control RIL-175 cells, overexpression of CCL17 delayed tumor progression; moreover, overexpression of IL-15 resulted in tumor regression in the Ras-driven CCA transplant model (**Fig. 4B**). We harvested subcutaneous tumors 13 days after transplantation, and measured tumor volume in control tumors (735.4±227.8 mm^3^), IL-15-overexpressed tumors (60.5±28.9 mm^3^), and CCL17-overexpressed tumors (295.7±141.5 mm^3^), again highlighting their critical role in delaying CCA tumor growth (**Fig. 4C and Fig. S8**). These results indicate that IL-15 and CCL17 suppress CCA tumor growth *in vivo*.

**Fig. 4.**
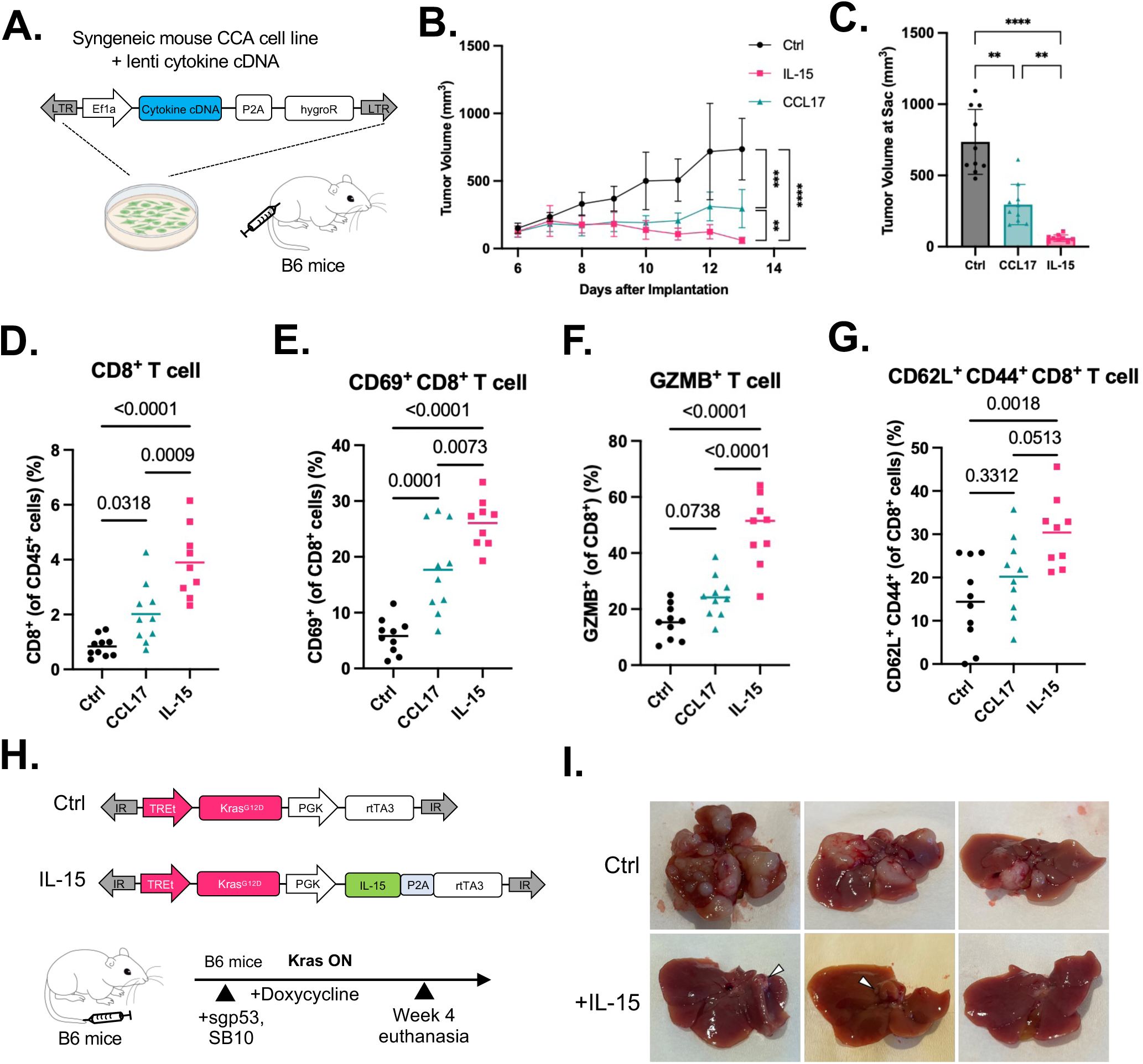
Murine IL-15 or CCL17 delays the progression of syngeneic mouse CCA xenograft tumors. **(A)** RIL-175 CCA cell lines from B6 genetic background were transduced with lentiviral vector encoding a constitutive Ef1α promoter and mouse cytokine/chemokine cDNAs or empty lentiviral vector as control. Xenograft model were made by injecting 10^6^ cells subcutaneously into immunocompetent B6 mice. **(B)** Overexpression of either IL-15 or CCL17 delayed tumor progression. Control, n=10; IL-15-O/E, n=9, CCL17-O/E, n=10. Data are presented as mean ± SD. **(C)** Tumor volume at day 13 after injection. Data are presented as mean ± SD. **(D-G)** Flow cytometry analysis in control, IL-15-overexpressed, and CCL17-overexpressed tumors at day 13 after injection. Lines represent mean value. **(E)** CD69, activated T cell marker. **(F)** GZMB, granzyme B. **(G)** CD62L^+^ CD44^+^, memory T cells markers. **(H)** IL-15 suppressed tumor formation in TKP CCA mouse model. IL-15-p2A was cloned into the TRE.*Kras* transposon plasmid and injected with transposase and sgp53-Cas9 plasmids into B6 mice. **(I)** Tumor burden with and without IL-15 expression were examined 4 weeks after injection. Arrows denote small tumors seen in the IL-15 group (n=3 mice). P-values were calculated by two-way ANOVA (B) or one-way ANOVA (C-G). \*\**P* <0.01, \*\*\**P* <0.001, \*\*\*\**P* <0.0001.

To test the anti-tumor effect of IL-15 in the liver, we cloned a new transposon construct constitutively expressing mouse IL-15: TRE-*Kras*^G12D^-PGK-*Il15*-rtTA (**Fig. 4H**). We co-delivered plasmids expressing SB transposon, transposase, and Cas9 nuclease with sgp53 into mouse liver by hydrodynamic tail-vein injection. In the TKP CCA model, IL-15 has a strong anti-tumor effect, demonstrated by a remarkable difference between surface tumor numbers of the IL-15 group and control group (**Fig. 4I**). These results provide further validation of anti-tumor effect and therapeutic potential of IL-15 in CCA.

### IL-15 and CCL17 recruits and activates CD8 T cells to delay tumor progression

To characterize immune cell infiltration and activation triggered by IL-15 or CCL17, we used flow cytometry to measure the immune cell types in subcutaneous tumors 13 days after transplantation in the RIL-175 CCA transplant model. Tumors that overexpress CCL17 or IL-15 triggered a greater influx of CD45^+^ immune cells (**Fig. S8A**). There was an increased infiltration of CD8^+^ T cells in IL-15-overexpressing CCA xenograft tumors, and to a lesser (but still significant) extent in CCL17-overexpressing tumors (**Fig. 4D**). Either IL-15 or CCL17 overexpression increased the level of activated CD8^+^ T cells (CD69-positive) (**Fig. 4E**). CD8^+^ T cells activated by IL-15 displayed elevated levels of tumor necrosis factor alpha (TNFα), granzyme B (GZMB), and interferon gamma (IFNγ) (**Fig. 4F and Fig. S9B,C**), indicating the activation of T lymphocytes. However, CCL17-activated CD8^+^ T cells only displayed increased expression level of TNFα (**Fig. 4F and Fig. S9B,C**). Moreover, CD8^+^ T lymphocytes activated by IL-15 exhibited a larger population of memory T cells, as assessed by staining of CD62L^+^ and CD44^+^ surface markers (**Fig. 4G**). These results suggest that IL-15 and CCL17 can recruit and activate CD8^+^ T lymphocytes that could contribute to CCA tumor regression.

### KRAS^G12D^ inhibitor MRTX1133 induces senescence and IL-15/Ccl17 expression in TKP ON cells

To test the pharmacological inhibition of Kras^G12D^, we treated TKP cells with a KRAS^G12D^ inhibitor MRTX1133(11). The inhibitor induces cellular senescence and reduces cell viability in a dose-dependent manner (**Fig. 5A-C**). MRTX1133 also upregulates IL-15 and Ccl17, as demonstrated by qRT-PCR (**Fig. 5D**), which is consistent with *Kras* withdrawal. Together, these results provide preclinical validation that KRAS^G12D^ inhibitor induces senescence and increases expression of IL-15 and Ccl17.

**Fig. 5.**
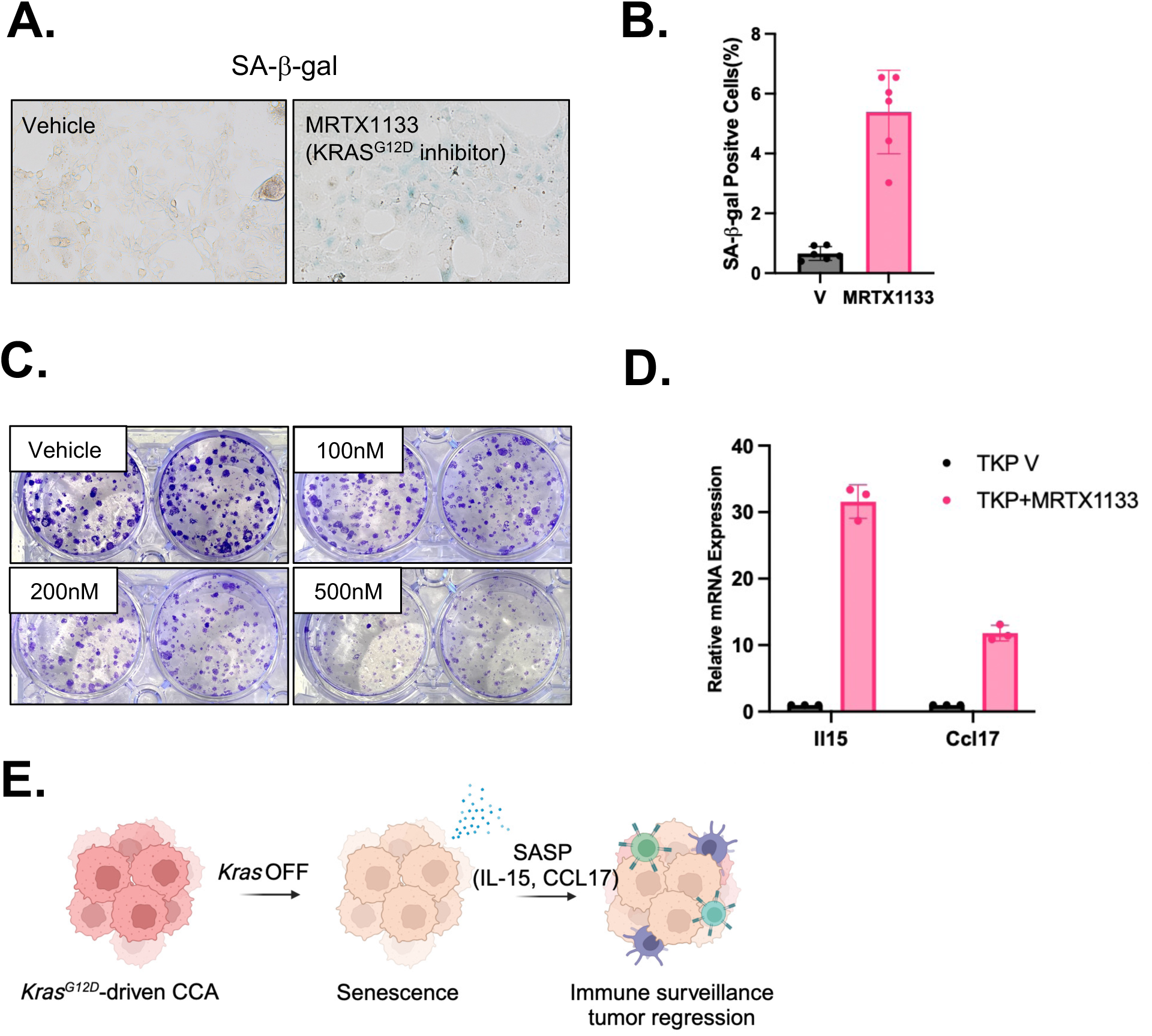
KRAS^G12D^ inhibitor MRTX1133 induces senescence and IL-15/CCL17 upregulation. **(A)** SA-β-gal staining of TKP cell line on Dox or on Dox + KRAS^G12D^ inhibitor MRTX1133 (72h). **(B)** Quantification of **(A)**. V, vehicle. Average is 6 of 10X fields. Data are presented as mean ± SD. **(C)** Clonogenic assay of TKP cells (*Kras* ON) treated with MRTX1133. **(D)** qRT-PCR analysis of IL-15 and CCL17 mRNA expression in TKP cell line treated with 500nM MRTX1133. Each group has three technical replicates. Data are presented as mean ± SD. **(E)** Model of CCA tumor regression induced by *Kras*^G12D^ withdrawal.

## Discussion

Our study establishes an *in vivo* model to study the mechanisms by which mutations in oncogenic *KRAS* contribute to CCA tumor maintenance. Conditional withdrawal of oncogenic *Kras* leads to significant immune-mediated tumor regression, highlighting the critical role of *Kras* in maintaining CCA tumors. Despite being driven by both oncogenic *Kras* and loss of *p53*, CCA tumors are sensitive to the withdrawal of *Kras^G12D^* even in the absence of tumor suppressor *p53*. This finding provides preclinical evidence for investigating KRAS-targeted therapies as a potential treatment for CCA, which usually progresses rapidly and is often diagnosed in an unresectable stage. This mouse model will be valuable in testing *Kras*^G12D^ inhibitors alone or in combination with immune-checkpoint therapies for CCA.

We propose that oncogenic *Kras* withdrawal results in cell cycle arrest through p53-independent senescence. Previous studies have demonstrated that *Kras^G12D^* withdrawal induces tumor regression in a lung adenocarcinoma mouse model and PDAC mouse models through apoptosis (18–20). By contrast, cellular senescence does not induce cell death but rather a stable, terminal arrest of proliferation (45). However, senescent cells can induce a non-cell autonomous program known as the SASP that can trigger immune surveillance (45). We proposed that oncogenic *Kras* withdrawal results in activation of TGF-β and SMAD signaling pathways, elevated expression of cell cycle inhibitors, and senescence. The senescent cells secrete proinflammatory factors, such as IL-15 and CCL17, to recruit and activate CD8^+^ T cells for tumor clearance (**Fig. 5E**). Consistently, recent studies showed that Kras^G12D^ inhibitor induces infiltration of neutrophils and CD8^+^ T cells in pancreatic tumor models (46–48). Our scRNA-Seq results revealed significant enrichment of T cells, dendritic cells, B cells, vascular cells, and CAFs in *Kras* OFF TKP tumor. Future studies will investigate the roles these cell subpopulations play in tumor regression following *Kras* withdrawal.

Our data demonstrate that both IL-15 and CCL17 can recruit and activate CD8^+^ T cells in CCA tumor, perhaps through different pathways. IL-15, but not CCL17, promotes T cell maturation into memory T cells. Also, both IL-15 and CCL17 did not trigger F4/80^+^ macrophage infiltration in xenograft CCA tumors (**Fig. S8D**). One potential drawback is that cytokine overexpression results in high levels of protein, which may not reflect normal physiological conditions. It nevertheless provides insight into cytokine/chemokine functions in CCA, which may help the prognosis process in clinics.

In summary, this study establishes a conditional CCA mouse model and provides functional evidence that oncogenic *Kras* withdrawal drives senescence and immune clearance, underscoring KRAS inhibition as a promising therapeutic strategy for CCA patients. Future work should test KRAS inhibitors in CCA mouse models to further validate KRAS targeted therapy as a potential therapeutic approach for CCA.

## Materials and Methods

### Sex as a biological variable

Both male and female C57BL/6 mice were involved in this study, and sex is not considered as a biological variable.

### Animals and TKP model

All animal protocols were approved by the UMass Chan Medical School Institutional Animal Care and Use Committee (IACUC). Mice were kept in specific pathogen-free conditions with access to food and water. The housing environment was controlled with a 12-hour light-dark cycle, with lights on from 07:00 to 19:00, a temperature range of 20–26°C, and humidity maintained between 30–70%. To make a TRE.KrasG12D/p53 knockout (TKP) CCA mouse model, 10 µg SB TREt-KrasG12D-PGK-rtTA3; 20 µg CMV-Cas9-sgp53(49); and 2µg CMV-SB10 transposase were delivered to 6-12-week-old C57BL/6 mice by hydrodynamic tail vein injection. To make a TKP model with a luciferase reporter, 15 µg SB TREt-Kras^G12D^-PGK-rtTA3; 20 µg CMV-Cas9-sgp53(49); and 6µg PT2-C-Luc-PGK-SB13 transposase (Addgene 20207) were delivered to 6-12-week-old C57BL/6 mice by hydrodynamic tail vein injection. Mouse strain C57BL/6 was purchased from Jackson Laboratory (Bar Harbor, Maine). Plasmid DNAs were prepared with EndoFree Maxiprep DNA Kit (Qiagen). Mice were fed on Doxycycline Diet 625 (Evigo) to induce the expression of Kras^G12D^. Four to five weeks after injection, mice were humanely euthanized by CO_2_ asphyxiation. Mouse livers were fixed in 10% (v/v) formalin overnight and dehydrated in 70% ethanol.

### Establishment of cell line from solid tumor

Fresh tumor tissues were obtained from TKP CCA mouse model (Kras ON) and minced into small pieces with autoclaved dissection tools. Tumors were then enzymatically dissociated in dispase medium containing 1.3 mg/mL dispase II (Invitrogen) and 20mM HEPE in Dulbecco’s Modification of Eagle’s Medium (DMEM) (Corning) for 45 minutes at 37°C. The resulting cell suspension was vortexed and centrifuged at 300 x g for 5 minutes. The pellets were then resuspended in fresh complete growth medium [DMEM with 10% FBS (vol/vol) and 1% penicillin-streptomycin (vol/vol)], plated in 10cm^2^ tissue culture dishes, and incubated at 37°C in 5% CO_2_ tissue culture incubator. Medium was changed every 2-3 days until cells reached confluence, after which they were passaged and expanded to establish the cell line. Mycoplasma testing was performed with MycoAlert Mycoplasma Detection Kit (Lonza) to ensure contamination-free cultures.

### Cell culture

Cells were cultured in DMEM, 10% serum (vol/vol) and 1% penicillin/streptomycin (vol/vol) under standard conditions, 37C in 5% CO_2_ tissue culture incubator. RIL-175 cell line is from Wen Xue(44). To induce Kras^G12D^ expression (Kras ON), TKP cell line was cultured in complete medium (DMEM+10%FBS+1%penicillin-streptomycin) with 1µg/mL doxycycline (Thermo Fisher). Kras OFF TKP cells were cultured with complete medium without doxycycline. All cell lines used were tested negative with MycoAlert Mycoplasma Detection Kit (Lonza).

### Colony formation Assay

TKP mouse liver cancer cells were seeded at 3,000 cells per well onto 6-well plates. Kras ON TKP cells were cultured in complete medium with 1µg/mL doxycycline, and Kras OFF cells were cultured in complete medium without doxycycline. After 8 days, cells were fixed with 0.5% glutaraldehyde, and stained with 0.5% crystal violet. The cells were then washed with water and dried at room temperature.

### Senescence-Associated Beta-Galactosidase Assay

SA-β-gal staining was conducted according to established protocols at pH 5.5 for mouse cells and tissues as described (38, 39, 50). Fresh frozen tumor tissue sections or cells grown in 6-well plates were fixed in 0.5% glutaraldehyde in PBS for 15 minutes. After fixation, the samples were rinsed with PBS containing 1 mM MgCl₂ and then incubated in a staining solution made up of PBS with 1 mM MgCl₂, 1 mg/mL X-Gal, and 5 mM of both potassium ferricyanide and potassium ferrocyanide for 4 to 18 hours. Tissue sections were counterstained with eosin.

### Histology and Immunohistochemistry

Dehydrated tissues were embedded in paraffin, cut into 4 μm sections, and stained with hematoxylin and eosin (H&E) by UMass Chan Medical School Morphology Core. For immunohistochemistry staining, tumor sections were deparaffinized with xylene and dehydrated with serial ethanol dilutions. To retrieve the antigen, tissues were boiled for 10 minutes with 1 mM citrate buffer at pH6.0 (Vector Labs). 3% hydrogen peroxide was used to block endogenous peroxidase activities for 10 mins at room temperature. Tissues were then blocked with 5% normal horse serum (Vector Labs) for 2 hours at room temperature and incubated with primary antibodies CK19 (EPNCIR127B; 1:400) (Abcam); CD8 (4SM15; 1:100) (Thermo); F4/80(D2S9R: 1:300) (Cell Signaling) 4C overnight. On the second day, tissues were incubated with secondary antibodies (HRP anti-rabbit IgG, HRP anti-rat IgG, Vector Labs) for 2 hours at room temperature, and diaminobenzidine (DAB) substrate/chromogen for development (Fisher Scientific). Slides were counterstained with hematoxylin and bluing reagent, dehydrated in ethanol and xylene, and sealed with a coverslip for long-term storage. H&E or IHC images were captured using a Leica DMi8 microscope. IHC slides were quantified by selection of 10 random fields per section, and positive cells per 20X field were counted.

### Immunofluorescence

Tissue sections were prepared for immunofluorescence staining using the standard protocol as described above for IHC. The following primary antibodies were used: CK19 (EPNCIR127B; 1:400) (Abcam); CD8 (4SM15; 1:100) (Thermo); CD69 (Cat# AF2386; 1:100) (R&D Systems). Primary antibodies were applied overnight. The following secondary antibodies were used: Alexa Flour anti-Goat 594 (Cat# A32758; 1:250); anti-Rat (Cat# A48272; 1:250); anti-Rabbit (Cat# A21207; 1:250). The secondary antibodies were applied 1 hour at room temperature. Fluorescence antibodies-labeled slides were mounted with Prolong Gold Antifade Mountant with DAPI (Cat# P36931; Thermo Fisher).

### Immunoblot Analysis

TKP cells were seeded in 6-well plate at density of 10^6^ and allowed to adhere overnight in complete growth medium with 1µg/mL doxycycline. Kras ON TKP cells were collected at 70-80% confluency, while Kras OFF TKP cells were collected at day 8 after doxycycline withdrawal (Kras OFF D8). TKP cells were starved before collection for immunoblot analysis. To induce starvation, complete growth medium was replaced with starvation medium (DMEM+0.1%FBS+1%penicillin-streptomycin) for 24 hours prior cell collection. Cells were lysed with RIPA buffer (Boston bioproducts) supplemented with protease inhibitor (Roche) and phosphatase inhibitor (Thermo Fisher). Cells were further lysed with sonication. Protein concentration was measured by BCA assay kit (Thermo Fisher). Equal amounts of protein (approximately 25µg) were loaded onto NuPAGE™ 4-12% Bis-Tris Protein Gels (Invitrogen) and run at 125 V for 90 mins. Protein was transferred to polyvinylidene fluoride (PVDF) membrane and incubated with indicated antibodies: anti-SMAD2 (5339T, CST), anti-phospho-SMAD2 (3108T, CST), anti-GAPDH (MAB374, EMD), followed by incubation with distinct fluorophore-conjugated secondary antibodies. Images were captured using the Odyssey system (Li-Cor Biosciences).

### RNA extraction and quantitative real-time PCR

Kras ON and Kras OFF D8 RNA was extracted with the RNeasy min kit (Qiagen). 500ng to 1 µg of RNA was used to synthesize complementary DNA (cDNA) using the high-capacity cDNA reverse-transcription kit (Thermo Fisher) according to the manufacturer’s protocol. Real-time quantitative PCR analysis was performed with SsoFast EvaGreen Supermix (Bio-rad). Beta-actin was served as an internal control for real-time PCR. All the primers for real-time PCR are listed in **Supporting Table S1**.

### Cytokine Array

TKP cells were plated in triplicate in 6-well plates. Conditioned media was collected from Kras ON and Kras OFF D8 TKP cells and diluted 1:2 with phosphate-buffered saline (PBS). RIL-175 Ras-driven CCA cell lines were plated in duplicates in 6-well plate. 48 hours later, Conditioned media was collected from RIL-175 cell lines and diluted 1:2 with PBS. Aliquots (100μL) of the conditioned media were analyzed using a multiplex immunoassay (Mouse Cytokine/Chemokine 44-Plex array) from Eve Technologies.

### RNA seq

RNA-Seq data were aligned to GRCm38 (mm10), and an RPKM matrix of transcript counts was generated. The R packages “limma” and “edgeR” were used to normalize the transcriptomic data and identify differentially expressed genes, respectively. Volcano plots were generated by R. The expression of senescence-associated cell cycle and SASP genes was from Ref (38). A heat map of gene expression (log2 fold change, LFC) was generated in R.

### GSEA analysis

Gene set enrichment analysis (GSEA) was performed using GSEA and R software (51). Ontology gene sets and immunologic signature gene sets were downloaded from the Human Molecular Signatures Database (52). The transcriptomic dataset used for GSEA was derived from RNA-seq of Kras ON and Kras OFF TKP cell lines. RNA sequencing data are available for download at the National Center for Biotechnology Information (GEO GSEXXXXX) or upon reasonable request from the authors.

### Single-Cell RNA Sequencing

Tumor tissues were dissected from the liver and dissociated on gentleMACS Tissue Dissociator (Miltenyi) using Tumor Dissociation Kit, Mouse (Miltenyi). The dissociated cells were filtered through a 30-μm strainer to remove debris, and red blood cells were lysed using RBC lysis buffer (Miltenyi). An aliquot of cell suspension was stained with Trypan blue, and viable cells were counted on the Cellometer Auto T4 Brightfield Cell Counter (Nexcelom). 10,000 viable cells were loaded onto the Chromium Controller (10x Genomics) using the Chromium Next GEM Single Cell 3’ Kit (10x Genomics). Library construction was performed following the manufacturer’s standard protocol (10x Genomics). The resulting libraries were sequenced on an Illumina NovaSeq X Plus platform.

Raw sequencing data were processed using Cell Ranger v7.0 (10x Genomics) (53) and aligned to the mouse genome (mm39), with the KrasG12D-rtTA3 transgene added to assemble the reference genome. Subsequent analysis was performed using Seurat v5.0 (54, 55). Quality control excluded cells with fewer than 200 detected genes or more than 10% mitochondrial gene content. The integrated dataset was normalized using the LogNormalize method. Datasets were integrated using Seurat’s CCA-based integration (55). The top 2,000 highly variable genes were identified by the Variance Stabilizing Transformation (VST) method (56) and used for principal component analysis (PCA). Cells were clustered using the Louvain algorithm (57) with a resolution of 0.5, based on the top 10 principal components, resulting in 15 subclusters across 13,329 cells. Clusters were visualized using Uniform Manifold Approximation and Projection (58).

Cell types were assigned to clusters based on the expression of known marker genes: immune cells (Ptprc), macrophages (Csf1r, C1qa, Cd68, Ccr2), neutrophils (S100a8, Cxcr2), T cells (Cd3d, Cd3e), vascular cells (Pecam1, Flt1), dendritic cells (Xcr1, Cd209a), tumor cells (Krt8, Krt18, rtTA3), cancer-associated fibroblasts (Dcn, Col1a1, Acta2), and B cells (Cd79a, Cd19, Igkc).

To assess the impact of KrasG12D withdrawal, the proportions of each cell type were calculated for Kras ON and Kras OFF tumors and compared using a Fisher’s exact test (FET) to determine enrichment. Within the T cell cluster, subpopulations were identified based on the expression of Cd8a (CD8+ T cells) and Cd69 (CD69+ T cells), including CD8+ CD69+ double-positive cells. The enrichment of these T cell subpopulations in Kras OFF versus Kras ON tumors was similarly evaluated using an FET.

### IL-15 and CCL17 overexpression with lentivirus

Murine *Il-15* and *Ccl17* cDNA was isolated from mouse cDNA library and cloned into a lentiviral vector with EF1α promotor. Lentiviruses were packaged by co-transfection of 293FS cells with expression constructs (*Il-15, Ccl17, or Empty*) envelope vectors (VSV-G), and packaging plasmids (Delta 8.2) using polybrene. RIL-175 cell lines were transduced with *Il-15*, *Ccl17* or control *Empty* constructs, and cell selection was performed with 500μg/mL hygromycin for 1-2 weeks. *Il-15* and *Ccl17* overexpression was confirmed by qRT-PCR and cytokine array.

### RIL-175 CCA Xenograft model

RIL-175 control, *Il-15* and *Ccl17* expressed cell lines were harvested for implantation when they reached 70-80% confluency. Prior to implantation, cells were trypsinized, resuspended in PBS, and mixed 1:1 ratio with Matrigel (Thermo Fisher) to a final volume of 200μL per injection. Six-week-old female C57BL/6 mice were purchased from Jackson Laboratory (Bar Harbor, ME). For xenograft establishment, mice were anesthetized with isoflurane (1-3% in oxygen) and injected subcutaneously into both the right flank and left flank with 200μL of the cell suspension (1 × 10^6^ cells). Tumor sizes were measured with digital calipers and calculated by the formula: 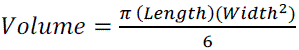 daily. RIL-175 xenograft tumors were harvested at day 13 after injection for further analysis.

### Flow cytometry analysis

To prepare single cell suspensions for flow cytometry analysis, RIL-175 xenograft tumors were minced with scissors into small pieces and placed in 5ml of collagenase buffer (1x HBSS with calcium and magnesium (Gibco), 1 mg/ml Collagenase V (Sigma-Aldrich), and 0.1 mg/ml DNase I) in C tubes and then processed using program 37C_m_TDK1_1 on a gentleMACS Octo dissociator with heaters (Miltenyi Biotec). Spleen was placed in 3 ml of PBS with 2% FBS in C tubes and dissociated using program m_spleen_01 on a gentleMACS Octo dissociator with heaters (Miltenyi Biotec). Dissociated tissue was passed through a 70 μm cell strainer and centrifuged at 500 x g for 5 minutes. To remove red blood cells, samples were lysed with ACK lysis buffer (Quality Biological) for 5 minutes and centrifuged and resuspended in PBS with 2% FBS. Then, samples were blocked with anti-CD16/32 (FC block, BD Pharmigen) for 20 minutes and then incubated with the following antibodies for 30 minutes on ice: CD45 AF700 (30-F11; 1:320), NK1.1 BV605 (PK136; 1:200), CD3 BV650 (17A2; 1:300), CD8 PE-Cy7 (53-6.7; 1:400), CD62L PE-Dazzle 594 (MEL-14; 1:200), CD44 BV785 (IM7; 1:100), CD69 APC-Cy7 (H1.2F3; 1:200), F4/80 APC (BM8; 1:200) (Biolegend). DAPI was used to distinguish live/dead cells. Flow cytometry was performed on an BD FACSymphony A5. Data was collected using BD FACSDiva Software, version 8.0 and analyzed using FlowJo, version 10.8.1 (TreeStar).

For analysis of Granzyme B (GZMB), Interferon-gamma (INFγ), and Tumor Necrosis Factor-alpha (TNFα) expression in NK and T cells, dissociated tumor tissue were resuspended in RPMI media with 10% FBS and 1% penicillin-streptomycin and incubated for 4 hours with PMA (20 ng/ml, Sigma-Aldrich), Ionomycin (1 μg/ml, STEMCELL technologies), and monensin (2μM, Biolegend) in a humidified incubator at 37°C with 5% CO2. Cell surface staining was first performed with CD45 AF700 (30-F11; 1:320), NK1.1 BV605 (PK136; 1:200), CD3 BV650 (17A2; 1:300), CD8 APC-Cy7 (53-6.7; 1:200) (Biolegend). For intracellular staining, samples were then fixed, permeabilized using the Foxp3/transcription factor staining buffer set (eBioscience), and then stained with GZMB APC (GB11, Biolegend; 1:100), INFγ V450 (XMG 1.2; 1:100), and TNFα PE-Cy7 (MP6-XT22; 1:100). GZMB, INFγ, and TNFα expression was evaluated by gating on CD3−NK1.1+ NK cells and CD3+CD8+ T cells on an BD LSR II flow cytometer as described above.

### IVIS Bioluminescent imaging

At the indicated time post injection, mice were given 200μl luciferin (Perkin Elmer) (15 mg/ml) intraperitoneally. Mice were then anesthetized with 3-5 minutes of 1-3% isoflurane. Fifteen minutes after luciferin injection, mice were loaded into the Perkin Elmer IVIS machine for capture of luminescent signal (1-minute standard exposure, scale 5e4 to 5e5 radiance)(59).

### Statistical Analysis

All statistical analyses were performed using GraphPad Prism 9.

## Acknowledgments

We thank D. Conte for editing the manuscript. We thank Y. Liu in the UMass Morphology Core for support, Chun-Qing Song for cloning the TRE.Kras plasmid, and G. Cottle for mouse injections. This work was supported by grants from the National Institutes of Health (R01CA275945, R01GM150279, P01HL158506, and UH3HL147367), Friedreich’s Ataxia Research Alliance (to W.X.) and grants from the National Cancer Institute (R00CA241110, R37CA282805) (to M.R).

---

**Fig. S1.**
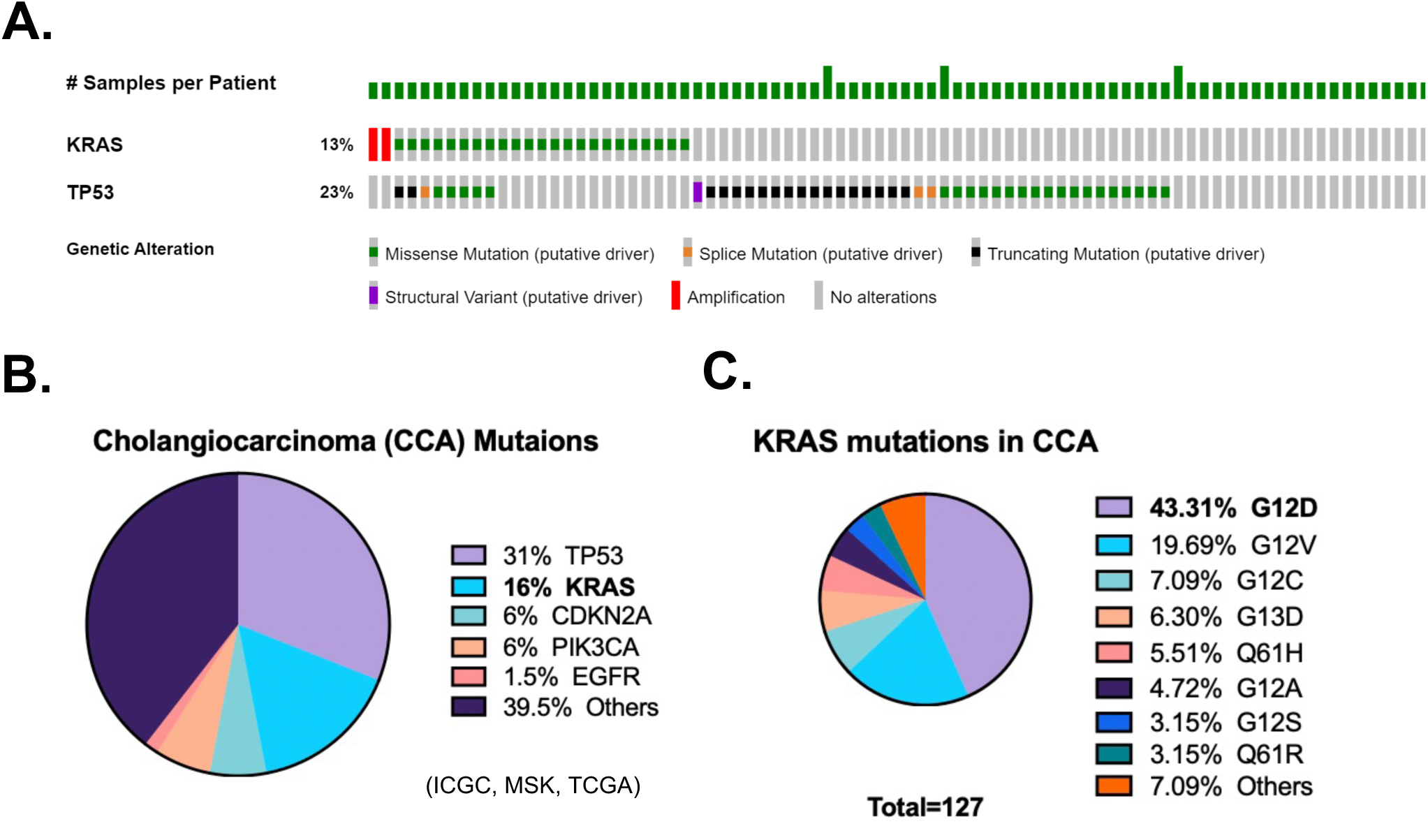
TRE.*Kras*^G12D^/*Trp53* knockout CCA mouse model. **(A)** Cancer genomics study of 195 CCA patients showed that Kras and p53 are frequently mutated in CCA. Data are from cBioPortal <u>Cholangiocarcinoma (MSK, Clin Cancer Res 2018)</u>. **(B)** Top genes mutated in CCA. **(C)** Distribution of Kras mutations (Zhou et al, JAMA Surg., 2021).

**Fig. S2.**
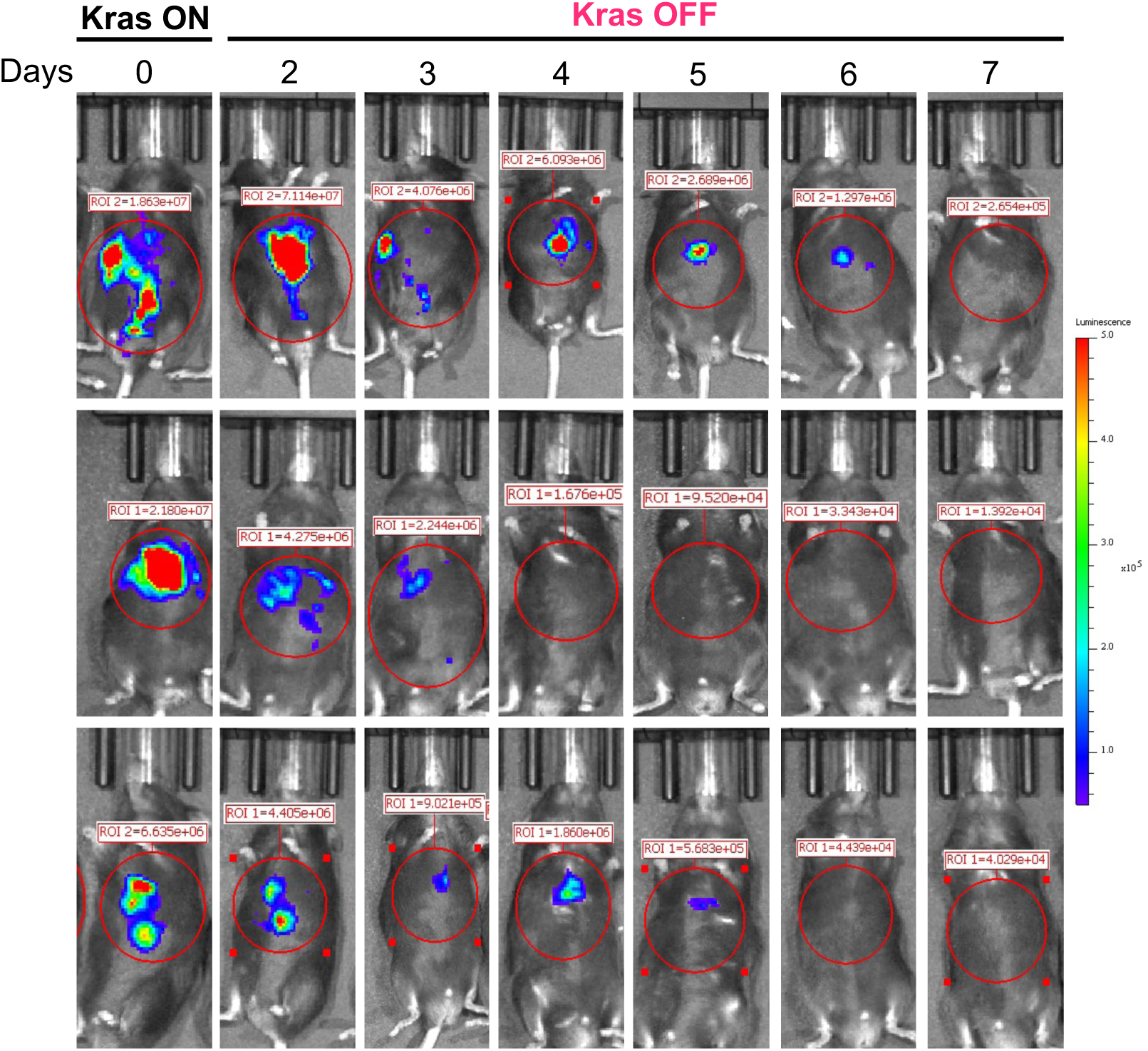
*Kras*^G12D^ withdrawal results in liver tumor regression in TKP model. Luciferase imaging shows that *Kras*^G12D^ withdrawal leads to tumor regression (n=3 mice). Luciferase radiance scale is 5e4-5e5.

**Fig. S3.**
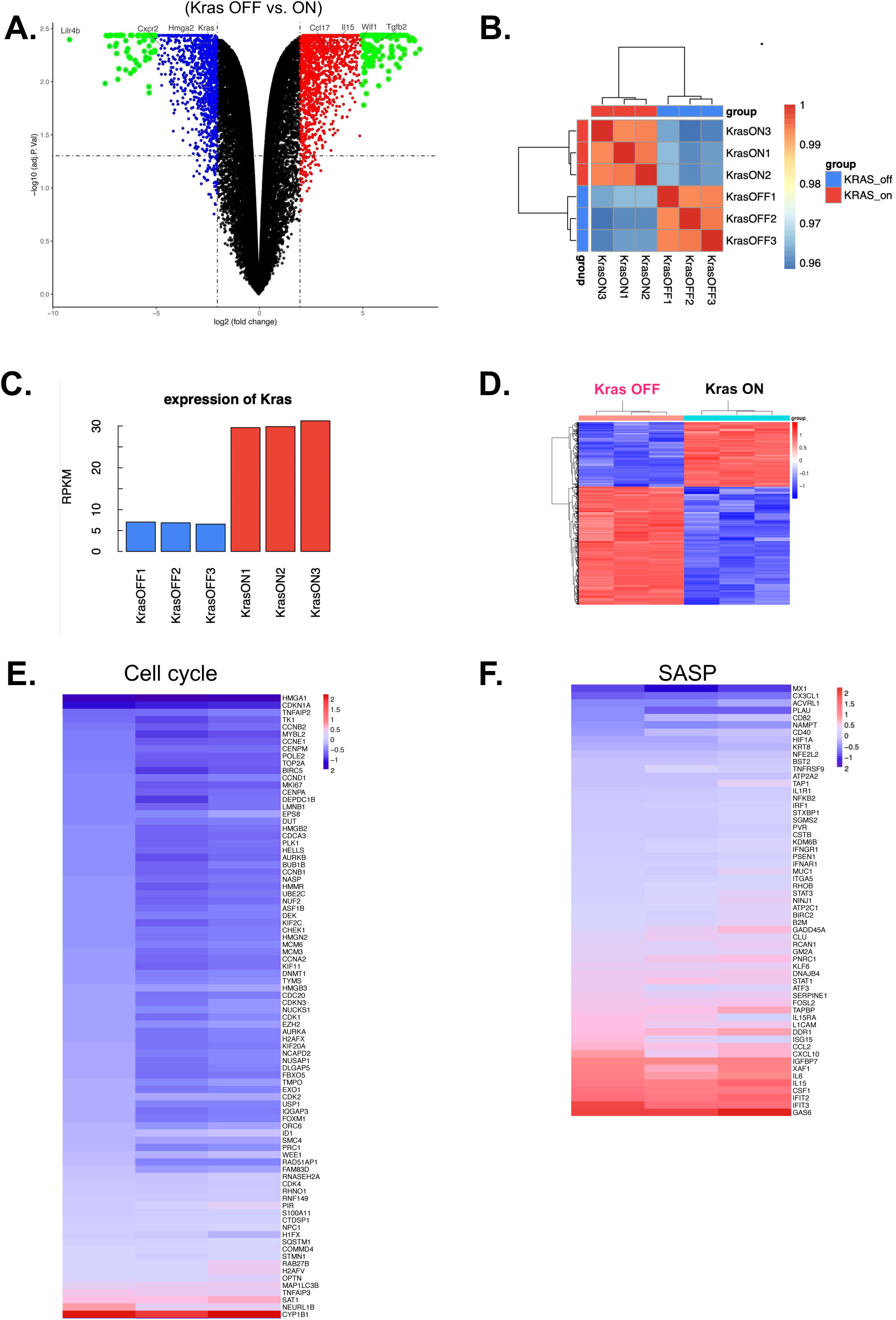
RNA-Seq analysis of *Kras* OFF vs. *Kras* ON TKP cells. **(A)** Volcano plot of differentially expressed genes comparing *Kras* OFF and *Kras* ON TKP cells. The x-axis represents log2 fold change of gene expression. Genes with a fold change greater than two were highlighted in blue (downregulated in *Kras* OFF vs. *Kras* ON) and red (upregulated in *Kras* OFF vs. *Kras* ON). Genes with a Log2 fold change greater than 5 were highlighted in green. The y-axis represents –log10 p-value. *Lilr4b*, Leukocyte Immunoglobulin-Like Receptor Subfamily B Member 4. *Cxcr2*, C-X-C Motif Chemokine Receptor 2. *Wif1*, WNT inhibitory factor 1. **(B)** Pearson correlation of gene expression between *Kras* ON and *Kras* OFF TKP cell lines. **(C)** RPKM showing difference in Kras expression between *Kras* OFF and *Kras* ON TKP cells. (**D)** Heatmap of differentially expressed transcripts log2-fold change in *Kras* OFF TKP cells compared with *Kras* ON cells (n=3 per group, genes that have log2-fold change greater than 2 were shown). **(E-F)** Heatmap of SASP and senescence-associated cell cycle gene expression. Three biological replicates were shown.

**Fig. S4.**
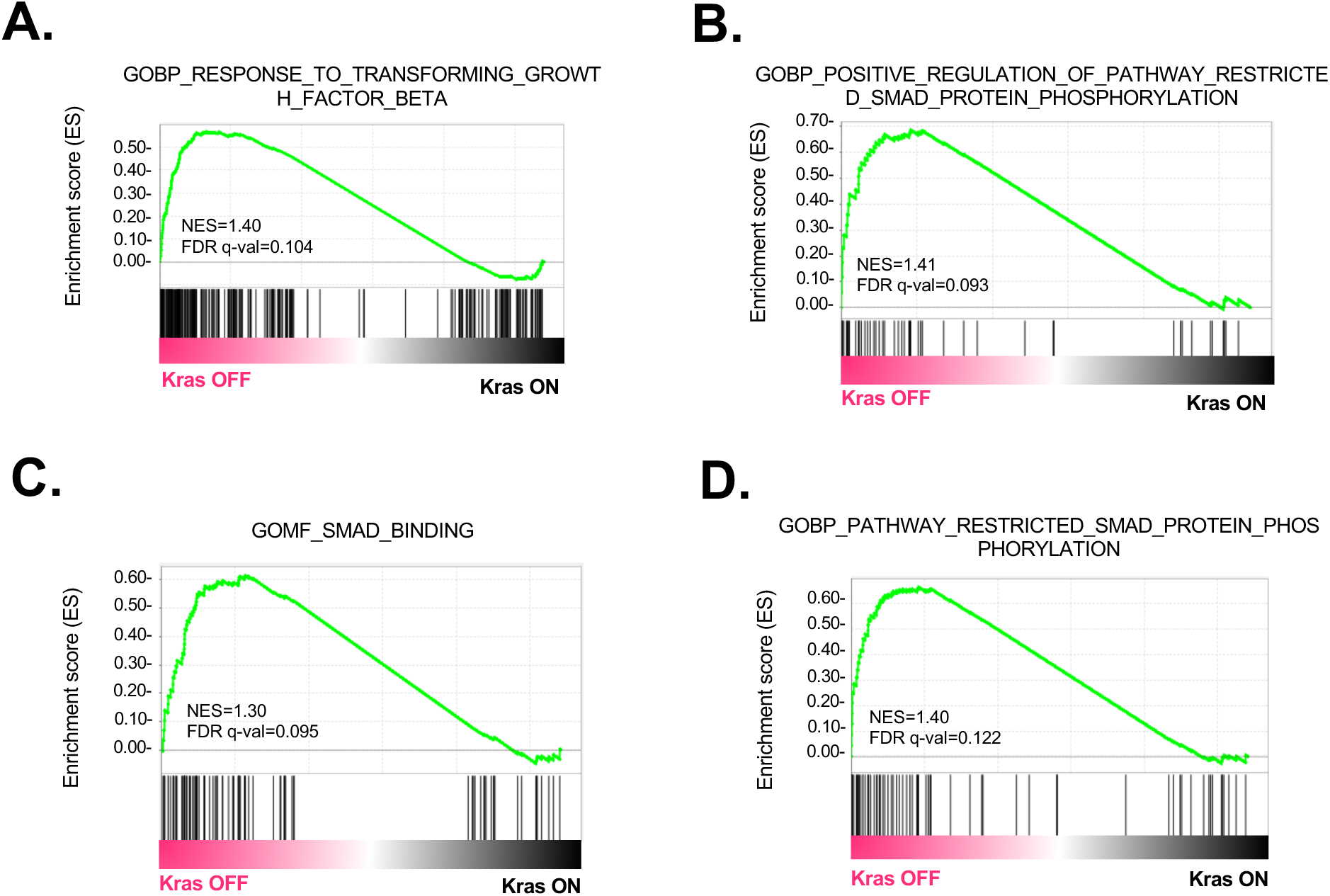
GSEA analysis of gene expression data comparing *Kras* ON and *Kras* OFF D8 TKP cell lines. (**A**) GSEA analysis of transforming growth factor beta pathway enrichment in *Kras* ON and *Kras* OFF cells. **(B-D)** GSEA analysis of SMAD signaling pathway enrichment in *Kras* ON and *Kras* OFF cells. NES, normalized enrichment score. FDR, false discovery rate.

**Fig. S5.**
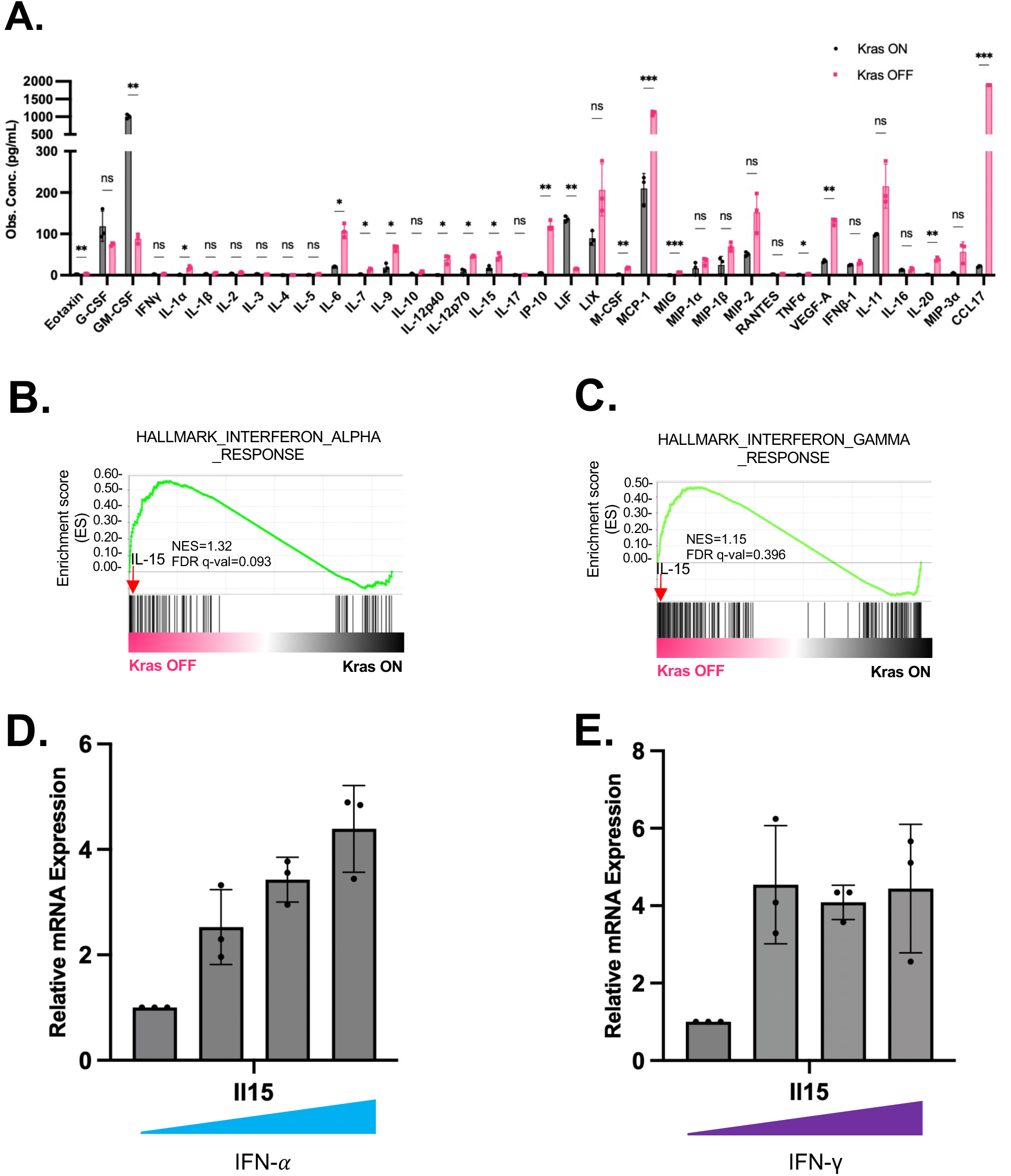
Senescence-associated secretory phenotype induced by *Kras*^G12D^ withdrawal. **(A)** Full list of cytokine array. Each group have three biological replicates. One outlier from Ccl17 group is omitted. Data are presented as mean ± SD. P-values were calculated by paired t-test. \**P* <0.05, \*\**P* <0.01, \*\*\**P* <0.001. **(B-C)** GSEA analysis of interferon alpha and gamma pathway enrichment in TKP *Kras* ON and TKP *Kras* OFF cells. IL-15 is the top 6^th^ and 4^th^ gene in the interferon alpha and gamma pathways, respectively. **(D-E)** qRT-PCR analysis of IL-15 expression in TKP cell line (on Dox) treated with increasing concentrations of IFN-alpha and IFN-gamma. Each group has three technical replicates. Data are presented as mean ± SD. IFN-alpha concentration used: 0, 10, 50, and 100ng/mL. IFN-gamma concentration used: 0, 100, 500, and 1000ng/mL.

**Fig. S6.**
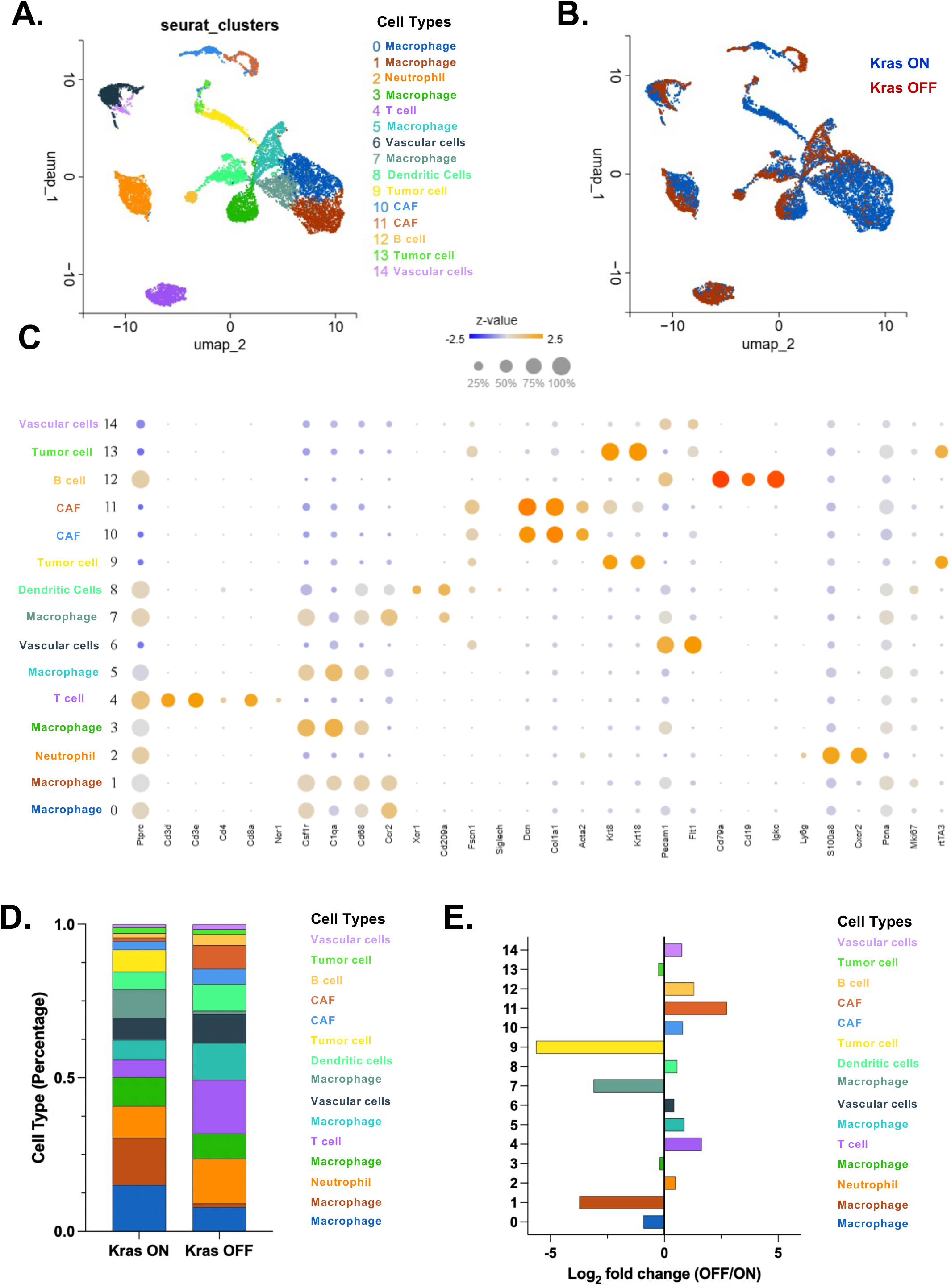
Single cell transcriptomic landscape reveals distinct cellular subpopulations in *Kras*^G12D^ ON and *Kras*^G12D^ OFF tumors. **(A)** UMAP visualization of cell clusters identified by scRNA-seq of 13,329 cells from one *Kras* ON and one *Kras* OFF tumors. **(B)** UMAP visualization of cell clusters color-coded for *Kras* ON and *Kras* OFF. **(C)** Dot plot showing marker gene for the cell subpopulations. **(D)** Bar plot showing proportions of cell subpopulations in *Kras* ON and *Kras* OFF tumors. **(E)** Log_2_ fold change in proportions of cell subpopulation between *Kras* OFF and *Kras* ON tumors.

**Fig. S6.**
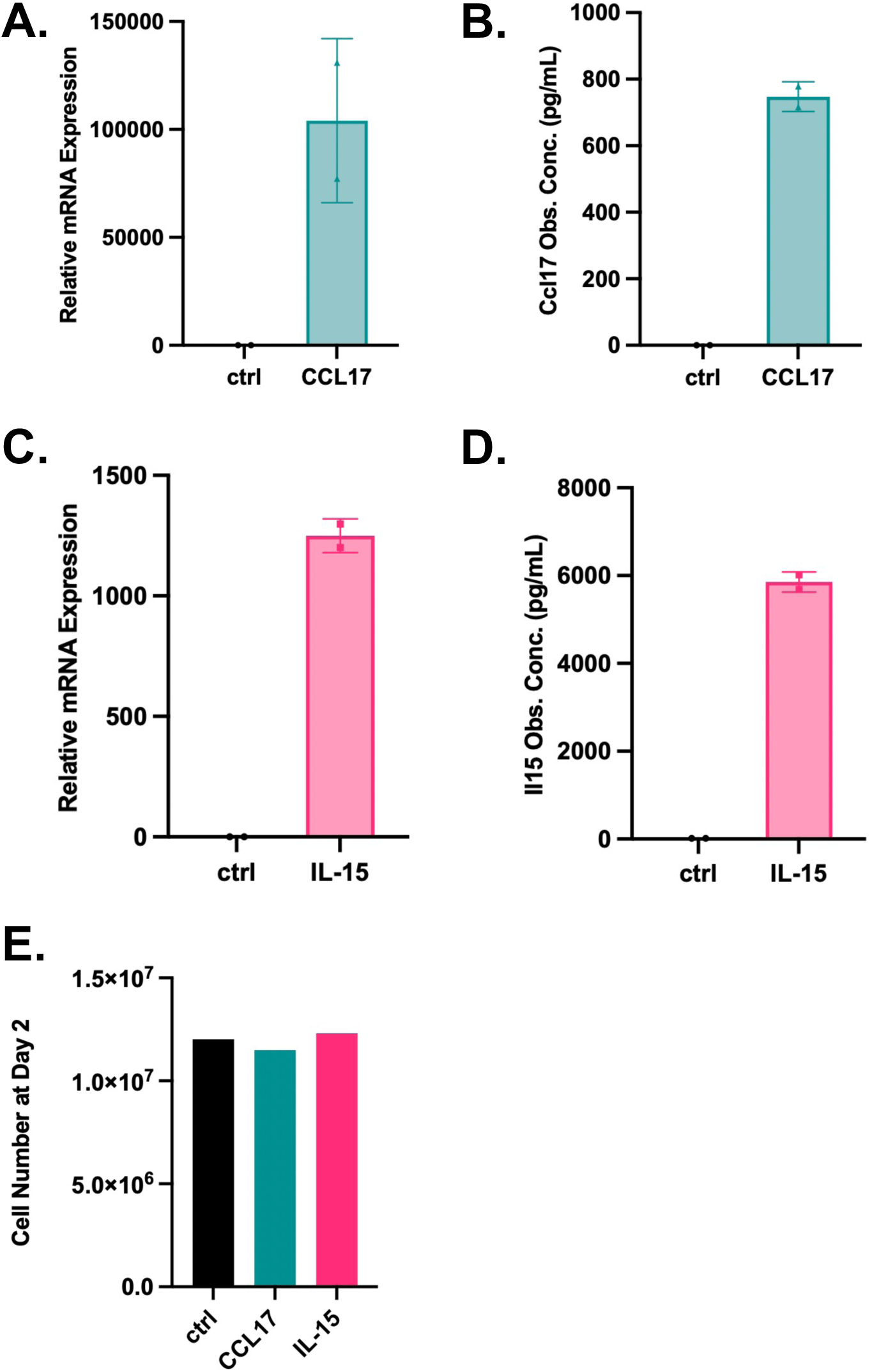
Overexpression of mouse *Il15* and *Ccl17* in RIL-175 Ras-driven CCA cell line. (**A-B**) qRT-PCR and cytokine array analysis of Ccl17 expression and secretion in RIL-175-Ccl17 compared to control. **(C-D)** qRT-PCR and cytokine array analysis of Il15 expression and secretion in RIL-175-Il15 compared to control. Data are presented as mean ± SD. **(E)** Cell number of control, Ccl17-expressed and Il15-expressed RIL-175 cell lines at day 2. Cells were seeded at 2.2*10^6^ at passage.

**Fig. S7.**
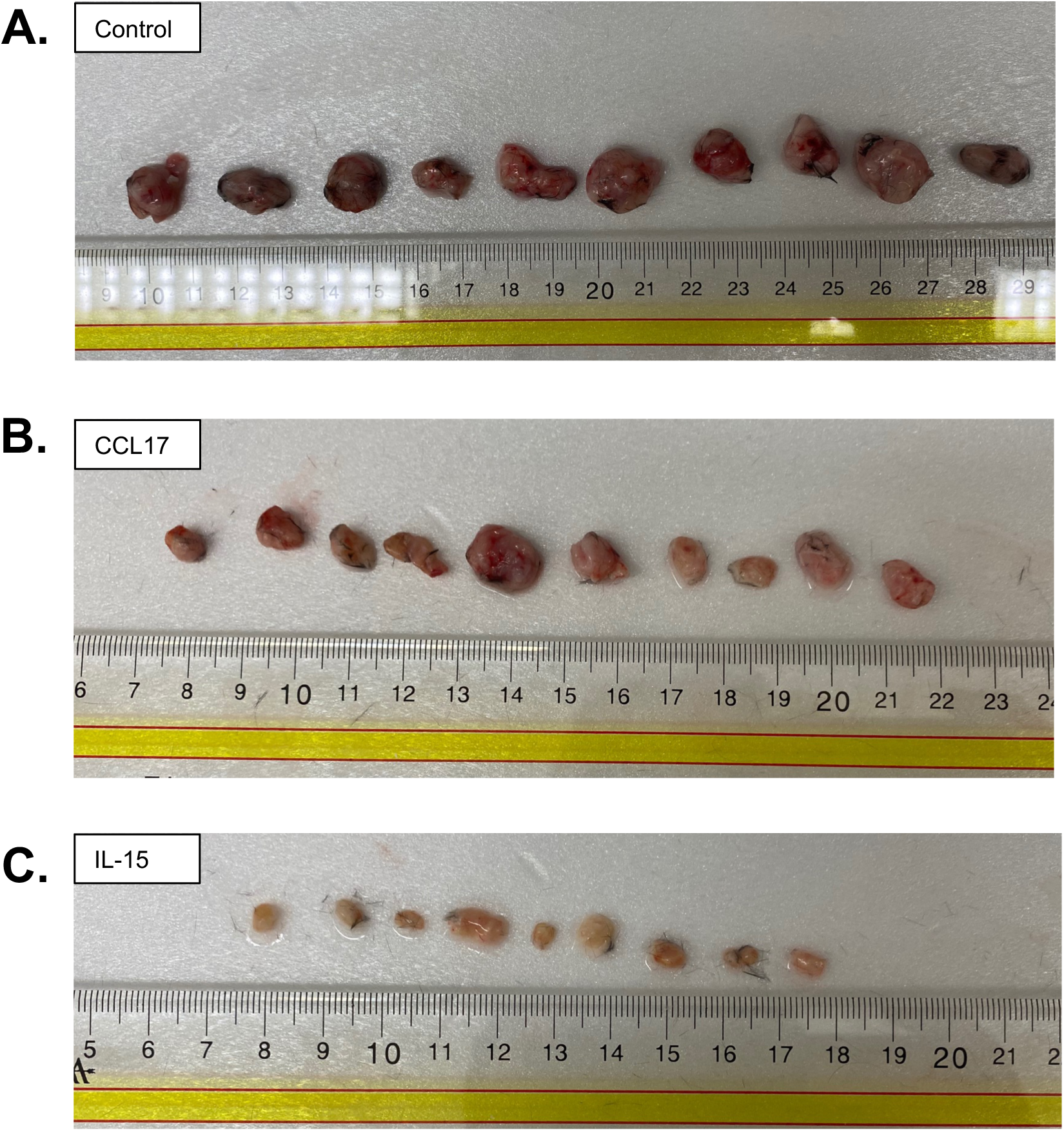
Gross tumor appearance of RIL-175 xenograft model at day 13 after implantation. Tumor appearance of **(A)** RIL-175 control, **(B)** CCL17, and **(C)** IL15 xenograft tumors. Ruler is in cm.

**Fig. S8.**
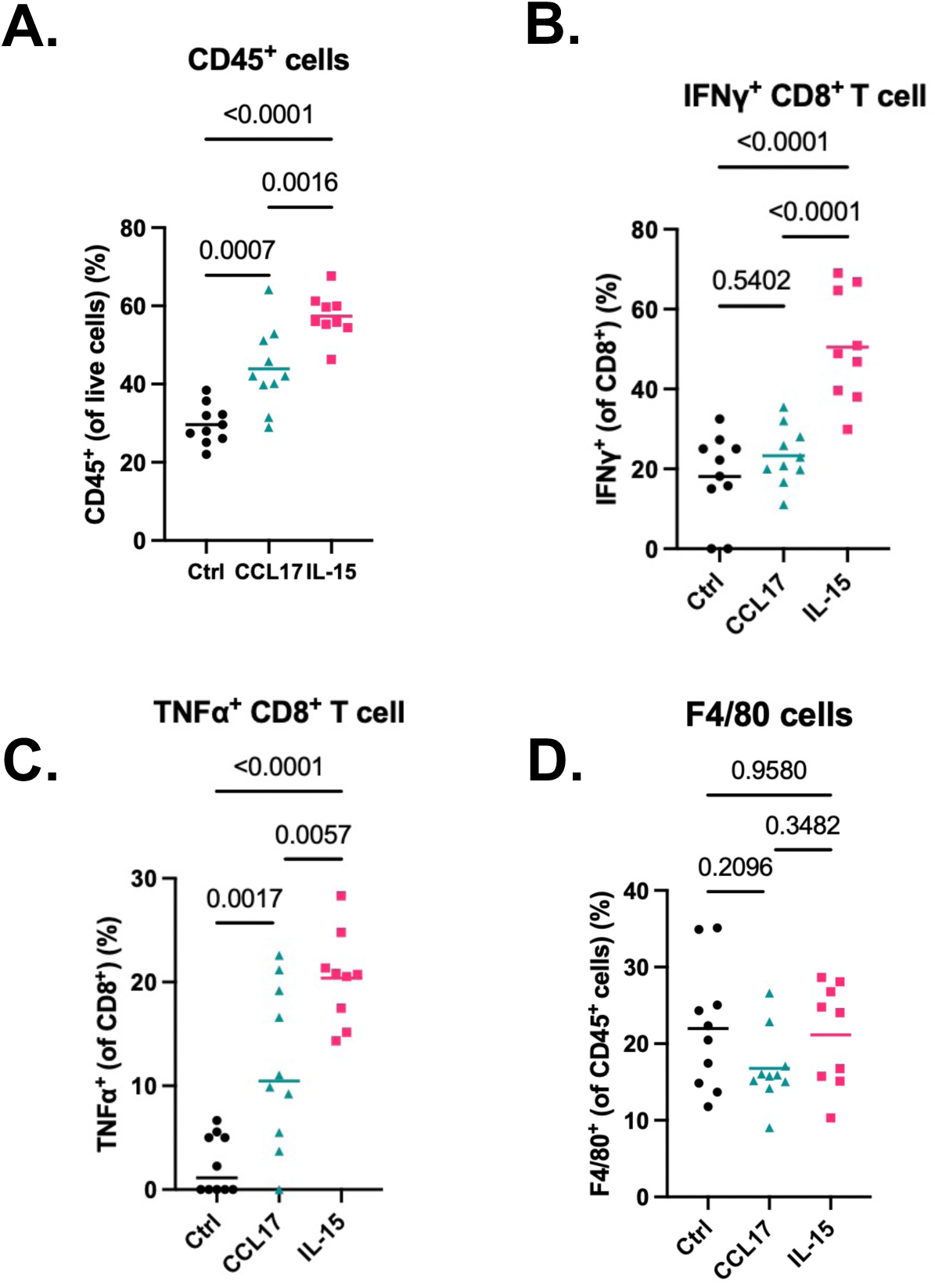
Flow cytometry analysis of RIL-175 xenograft tumors at day 13 after implantation. **(A)** CD45 positive cells. **(B-C)** IFNγ or TNFa positive cells within CD8 positive cell population. **(D)** F4/80 positive cells within CD45 positive cell population. P-values were calculated by one-way ANOVA.

