## Supplemental Figures for "KRAS withdrawal in Cholangiocarcinoma leads to immune infiltration and tumor regression"

### Fig.S1

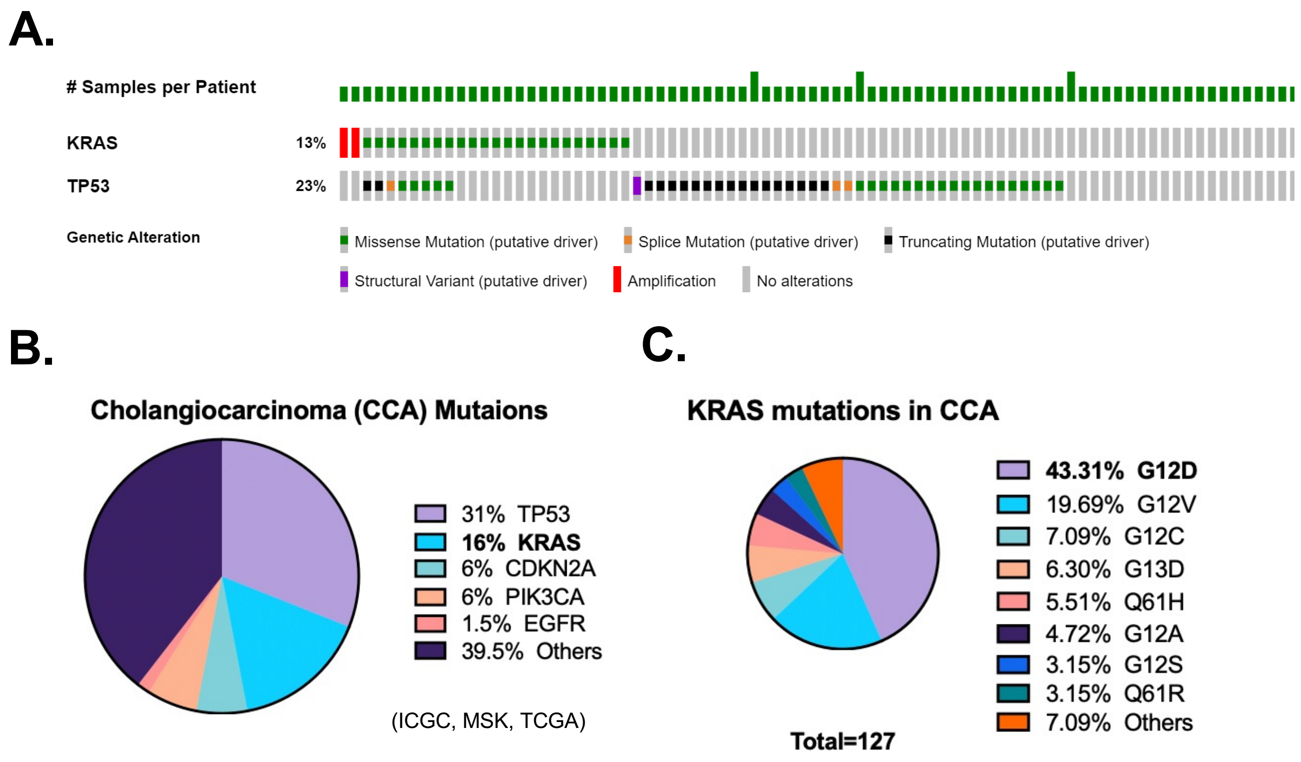


**Fig. S1. TRE.*Kras*$^{G12D}$/*Trp53* knockout CCA mouse model. (A)** Cancer genomics study of 195 CCA patients showed that Kras and p53 are frequently mutated in CCA. Data are from cBioPortal Cholangiocarcinoma (MSK, Clin Cancer Res 2018). **(B)** Top genes mutated in CCA. **(C)** Distribution of Kras mutations (Zhou et al, JAMA Surg., 2021).

### Fig.S2

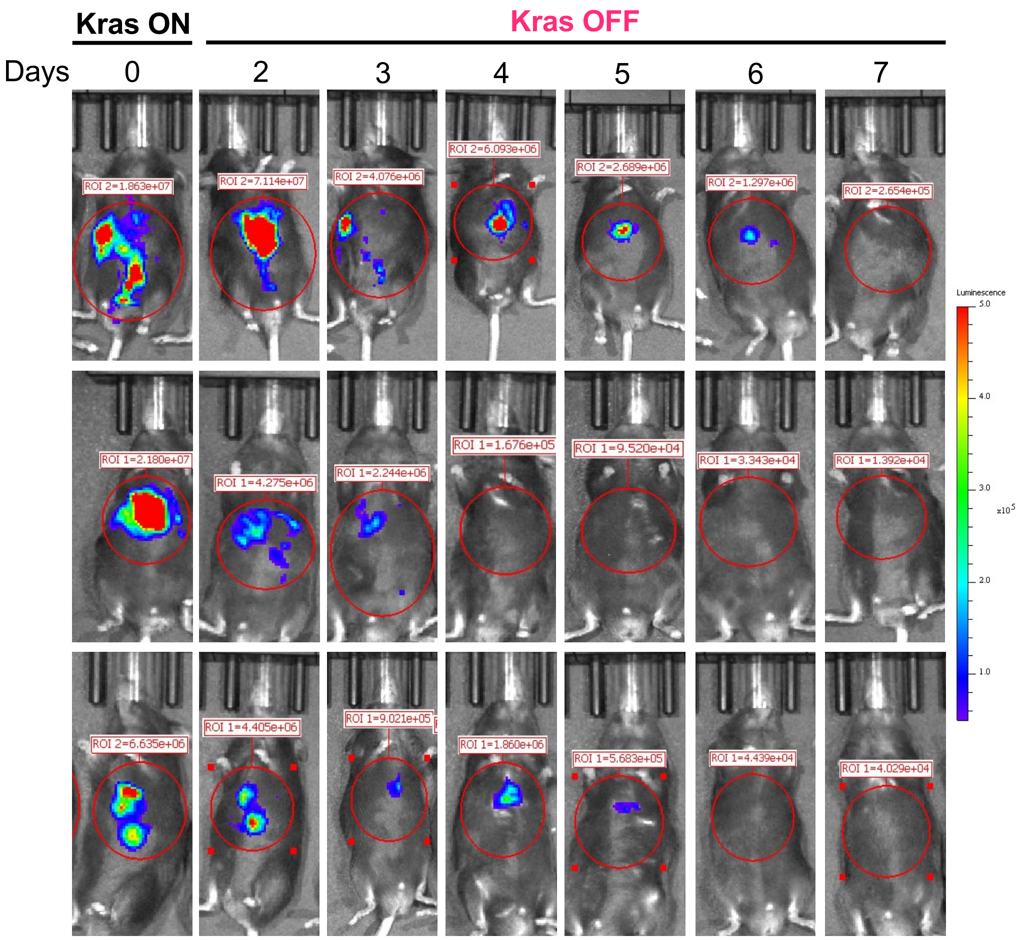


**Fig. S2. *Kras*$^{G12D}$ withdrawal results in liver tumor regression in TKP model.** Luciferase imaging shows that *Kras*$^{G12D}$ withdrawal leads to tumor regression (n=3 mice). Luciferase radiance scale is 5e4-5e5.

**Fig. S3.**

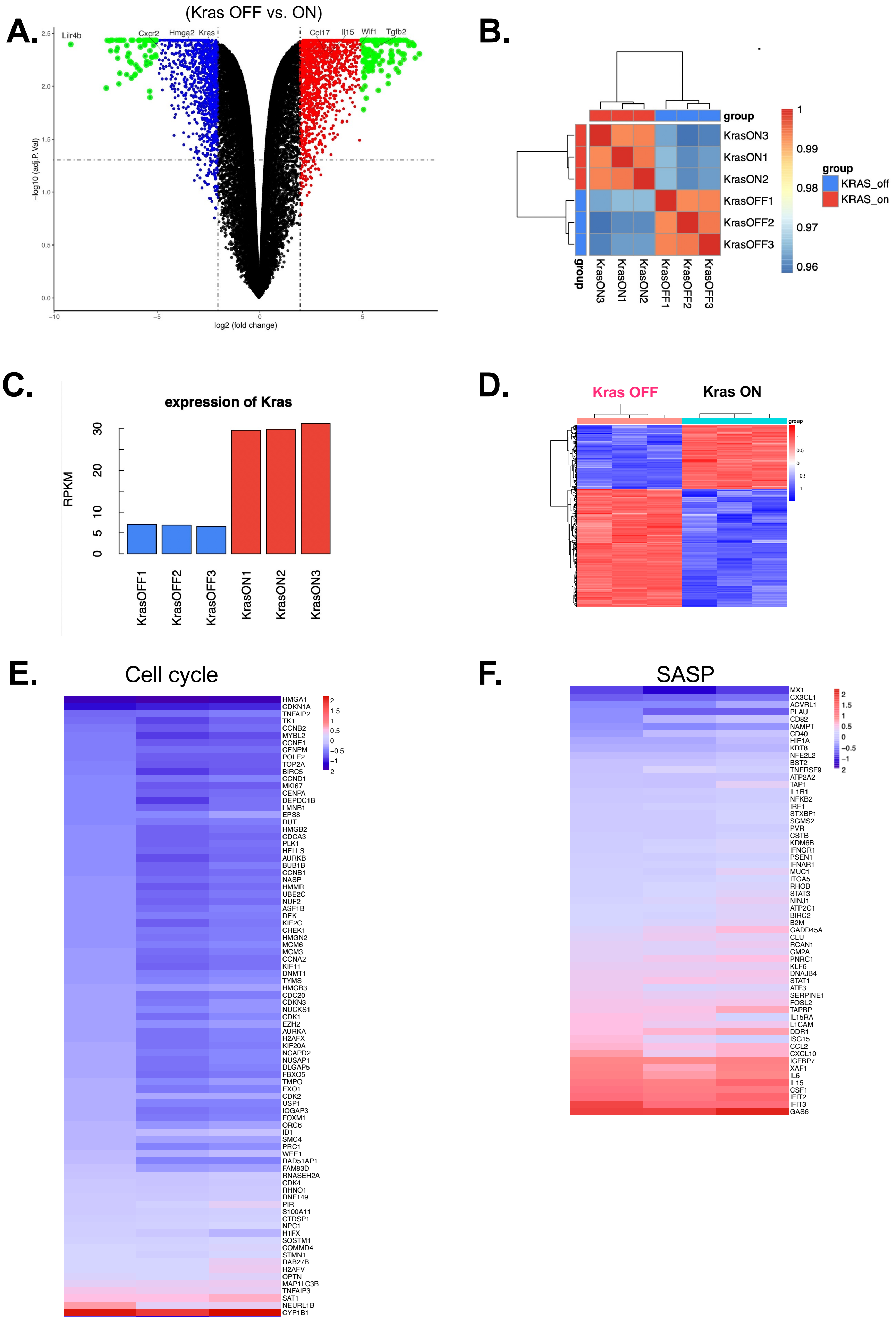
A.
(Kras OFF vs. ON)
Lilr4b
Cxcr2
Hmga2
Kras
Ccl17
Il15
Wif1
Tgfb2
−log10 (adj.P.Val)
log2 (fold change)
B.
group
KrasON3
KrasON1
KrasON2
KrasOFF1
KrasOFF2
KrasOFF3
group
KRAS_off
KRAS_on
C.
expression of Kras
RPKM
KrasOFF1
KrasOFF2
KrasOFF3
KrasON1
KrasON2
KrasON3
D.
Kras OFF
Kras ON
E.
Cell cycle
HMGA1
CDKN1A
TNFAIP2
TK1
CCNB2
MYBL2
CCNE1
CENPM
POLE2
TOP2A
BIRC5
CCND1
MKI67
CENPA
DEPDC1B
LMNB1
EPS8
DUT
HMGB2
CDCA3
PLK1
HELLS
AURKB
BUB1B
CCNB1
NASP
HMMR
UBE2C
NUF2
ASF1B
DEK
KIF2C
CHEK1
HMGN2
MCM6
MCM3
CCNA2
KIF11
DNMT1
TYMS
HMGB3
CDC20
CDKN3
NUCKS1
CDK1
EZH2
AURKA
H2AFX
KIF20A
NCAPD2
NUSAP1
DLGAP5
FBXO5
TMPO
EXO1
CDK2
USP1
IQGAP3
FOXM1
ORC6
ID1
SMC4
PRC1
WEE1
RAD51AP1
FAM83D
RNASEH2A
CDK4
RHNO1
RNF149
PIR
S100A11
CTDSP1
NPC1
H1FX
SQSTM1
COMMD4
STMN1
RAB27B
H2AFV
OPTN
MAP1LC3B
TNFAIP3
SAT1
NEURL1B
CYP1B1
F.
SASP
MX1
CX3CL1
ACVRL1
PLAU
CD82
NAMPT
CD40
HIF1A
KRT8
NFE2L2
BST2
TNFRSF9
ATP2A2
TAP1
IL1R1
NFKB2
IRF1
STXBP1
SGMS2
PVR
CSTB
KDM6B
IFNGR1
PSEN1
IFNAR1
MUC1
ITGA5
RHOB
STAT3
NINJ1
ATP2C1
BIRC2
B2M
GADD45A
CLU
RCAN1
GM2A
PNRC1
KLF6
DNAJB4
STAT1
ATF3
SERPINE1
FOSL2
TAPBP
IL15RA
L1CAM
DDR1
ISG15
CCL2
CXCL10
IGFBP7
XAF1
IL6
IL15
CSF1
IFIT2
IFIT3
GAS6

**Fig. S3. RNA-Seq analysis of *Kras* OFF vs. *Kras* ON TKP cells. (A)** Volcano plot of differentially expressed genes comparing *Kras* OFF and *Kras* ON TKP cells. The x-axis represents log2 fold change of gene expression. Genes with a fold change greater than two were highlighted in blue (downregulated in *Kras* OFF vs. *Kras* ON) and red (upregulated in *Kras* OFF vs. *Kras* ON). Genes with a Log2 fold change greater than 5 were highlighted in green. The y-axis represents –log10 p-value. *Lilr4b*, Leukocyte Immunoglobulin-Like Receptor Subfamily B Member 4. *Cxcr2*, C-X-C Motif Chemokine Receptor 2. *Wif1*, WNT inhibitory factor 1. **(B)** Pearson correlation of gene expression between *Kras* ON and *Kras* OFF TKP cell lines. **(C)** RPKM showing difference in Kras expression between *Kras* OFF and *Kras* ON TKP cells. (**D)** Heatmap of differentially expressed transcripts log2-fold change in *Kras* OFF TKP cells compared with *Kras* ON cells (n=3 per group, genes that have log2-fold change greater than 2 were shown). **(E-F)** Heatmap of SASP and senescence-associated cell cycle gene expression. Three biological replicates were shown.

### Fig.S4 GSEA

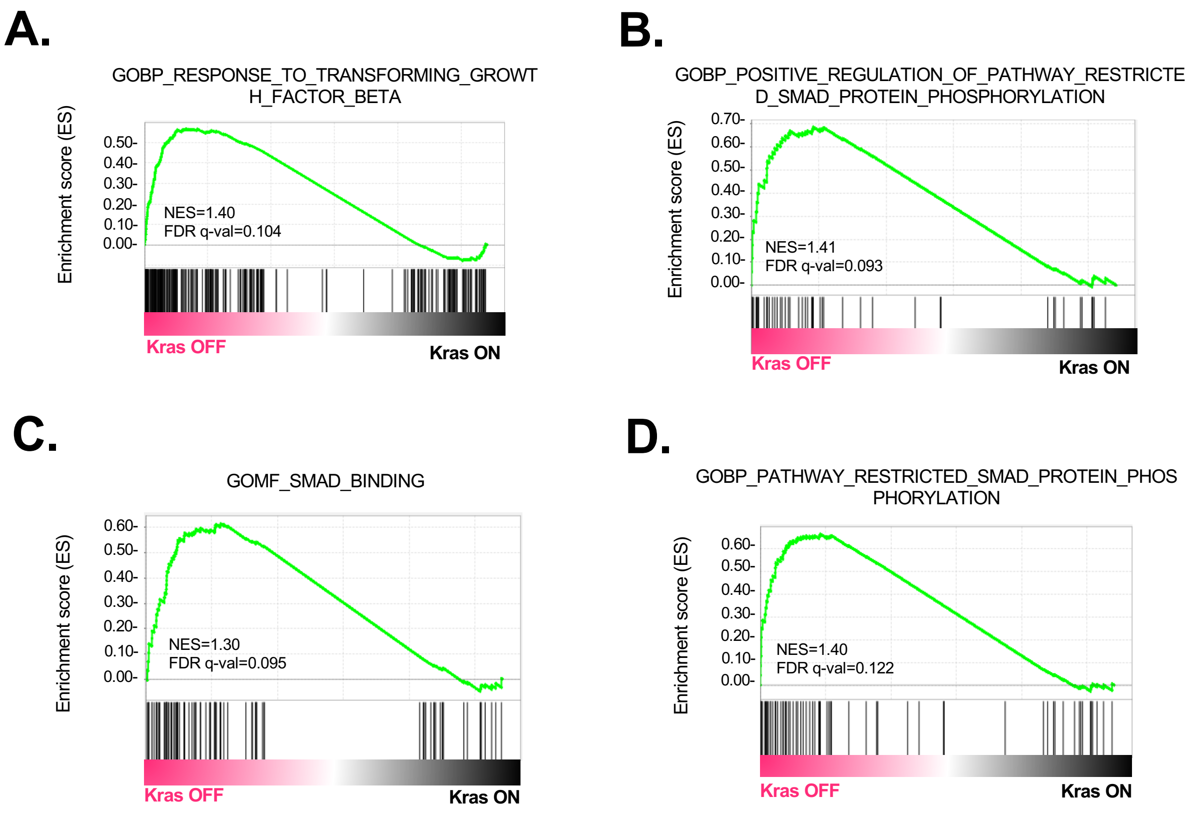


**Fig. S4. GSEA analysis of gene expression data comparing *Kras* ON and *Kras* OFF D8 TKP cell lines**. **(A)** GSEA analysis of transforming growth factor beta pathway enrichment in *Kras* ON and *Kras* OFF cells. **(B-D)** GSEA analysis of SMAD signaling pathway enrichment in *Kras* ON and *Kras* OFF cells. NES, normalized enrichment score. FDR, false discovery rate.

### Fig. S5

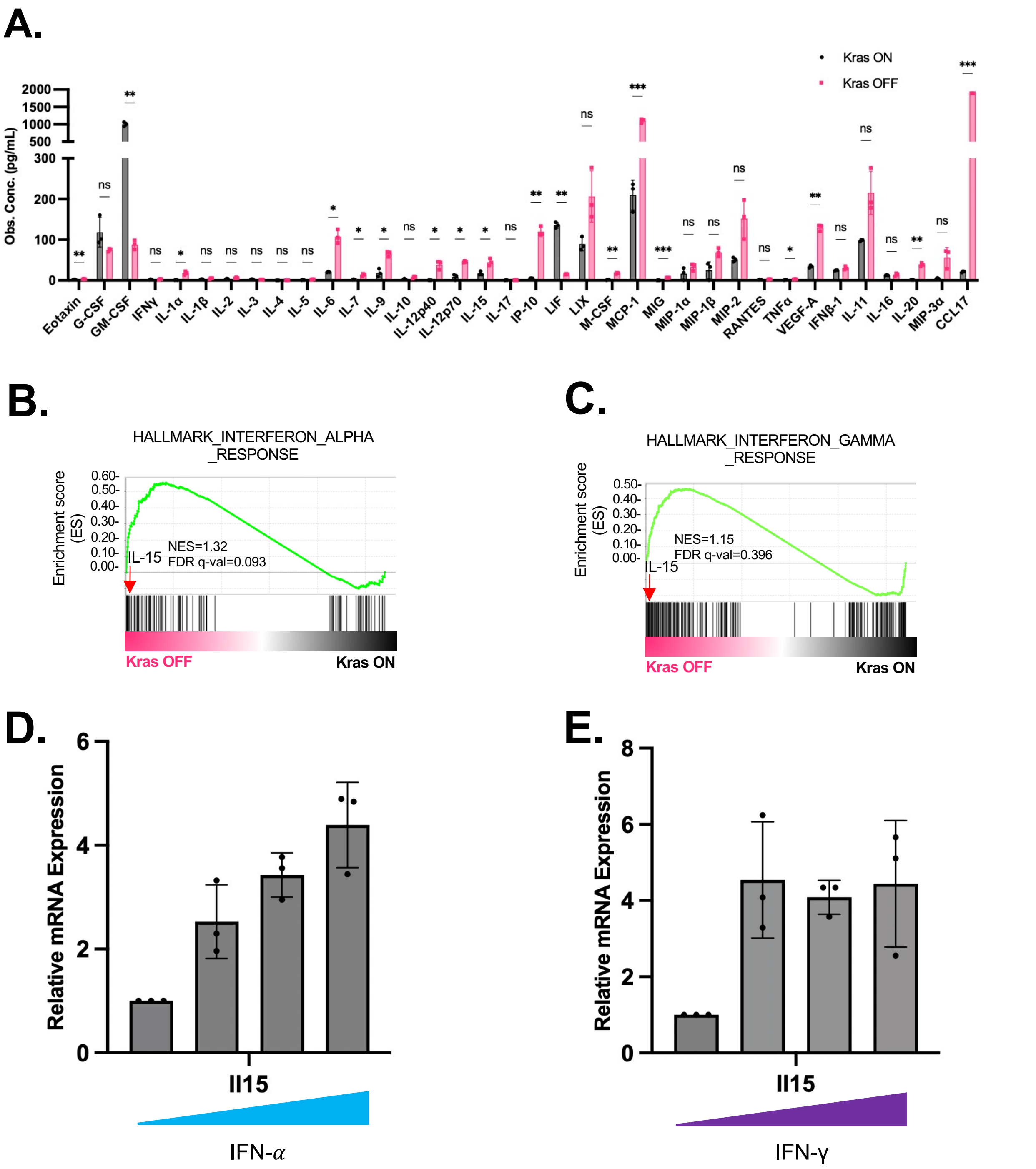


**Fig. S5. Senescence-associated secretory phenotype induced by *Kras*$^{G12D}$ withdrawal. (A)** Full list of cytokine array. Each group have three biological replicates. One outlier from Ccl17 group is omitted. Data are presented as mean ± SD. P-values were calculated by paired t-test. **P* <0.05, ***P* <0.01, ****P* <0.001. **(B-C)** GSEA analysis of interferon alpha and gamma pathway enrichment in TKP *Kras* ON and TKP *Kras* OFF cells. IL-15 is the top 6$^{th}$ and 4$^{th}$ gene in the interferon alpha and gamma pathways, respectively. **(D-E)** qRT-PCR analysis of IL-15 expression in TKP cell line (on Dox) treated with increasing concentrations of IFN-alpha and IFN-gamma. Each group has three technical replicates. Data are presented as mean ± SD. IFN-alpha concentration used: 0, 10, 50, and 100ng/mL . IFN-gamma concentration used: 0, 100, 500, and 1000ng/mL.

### Fig.S6

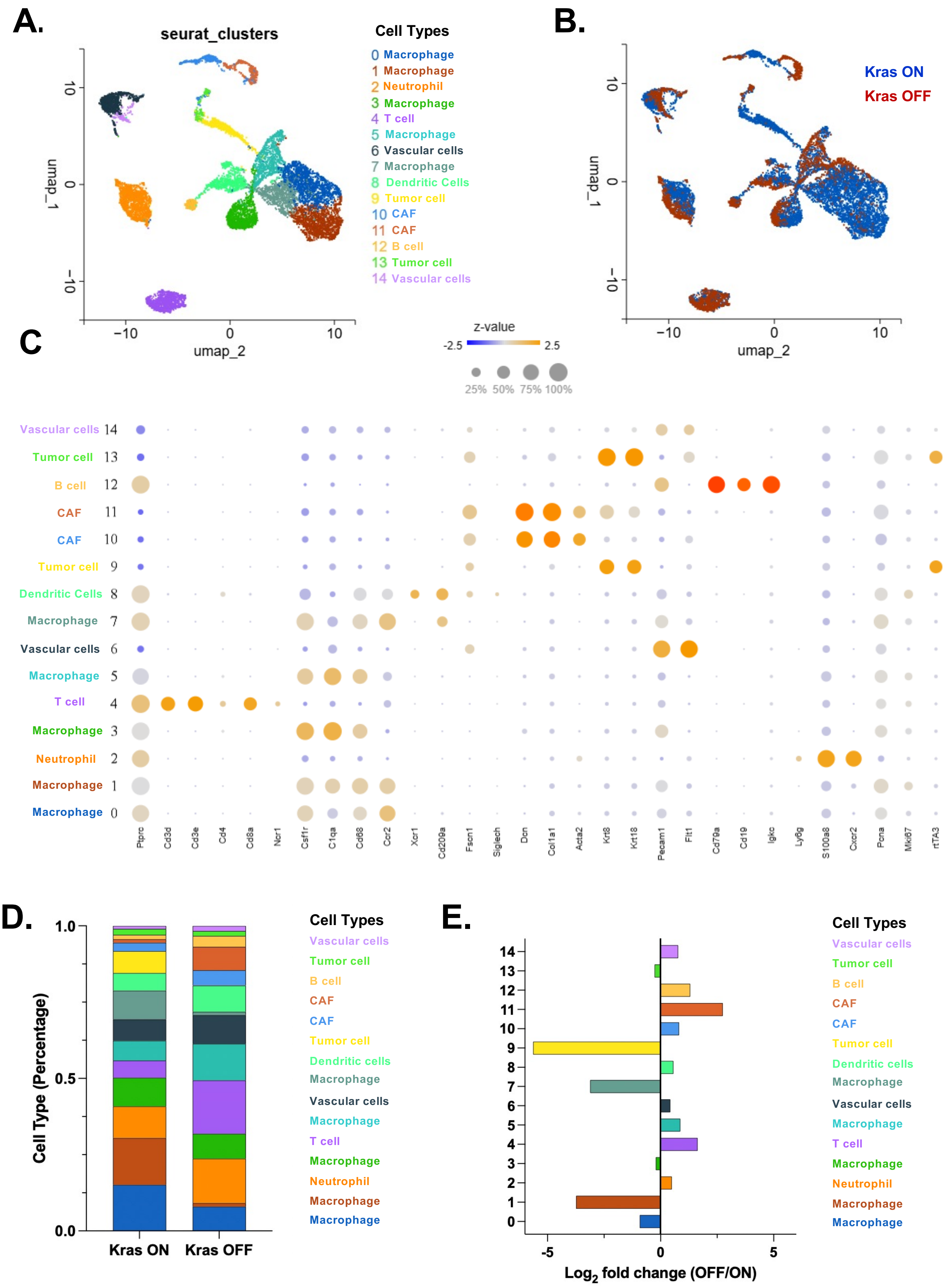
A.
seurat_clusters
Cell Types
0 Macrophage
1 Macrophage
2 Neutrophil
3 Macrophage
4 T cell
5 Macrophage
6 Vascular cells
7 Macrophage
8 Dendritic Cells
9 Tumor cell
10 CAF
11 CAF
12 B cell
13 Tumor cell
14 Vascular cells
umap_1
umap_2
B.
Kras ON
Kras OFF
C
z-value
-2.5
2.5
25% 50% 75% 100%
Ptprc Cd3d Cd3e Cd4 Cd8a Ncr1 Csf1r C1qa Cd68 Ccr2 Xcr1 Cd209a Fscn1 Siglech Dcn Col1a1 Acta2 Krt8 Krt18 Pecam1 Flt1 Cd79a Cd19 Igkc Ly6g S100a8 Cxcr2 Pcna Mki67 rtTA3
D.
Cell Type (Percentage)
Kras ON
Kras OFF
Cell Types
Vascular cells
Tumor cell
B cell
CAF
CAF
Tumor cell
Dendritic cells
Macrophage
Vascular cells
Macrophage
T cell
Macrophage
Neutrophil
Macrophage
Macrophage
E.
Log2 fold change (OFF/ON)
Cell Types

**Fig. S6. Single cell transcriptomic landscape reveals distinct cellular subpopulations in *Kras*$^{G12D}$ ON and *Kras*$^{G12D}$ OFF tumors. (A)** UMAP visualization of cell clusters identified by scRNA-seq of 13,329 cells from one *Kras* ON and one *Kras* OFF tumors. **(B)** UMAP visualization of cell clusters color-coded for *Kras* ON and *Kras* OFF. **(C)** Dot plot showing marker gene for the cell subpopulations. **(D)** Bar plot showing proportions of cell subpopulations in *Kras* ON and *Kras* OFF tumors. **(E)** $Log_2$ fold change in proportions of cell subpopulation between *Kras* OFF and *Kras* ON tumors.

Fig.S7

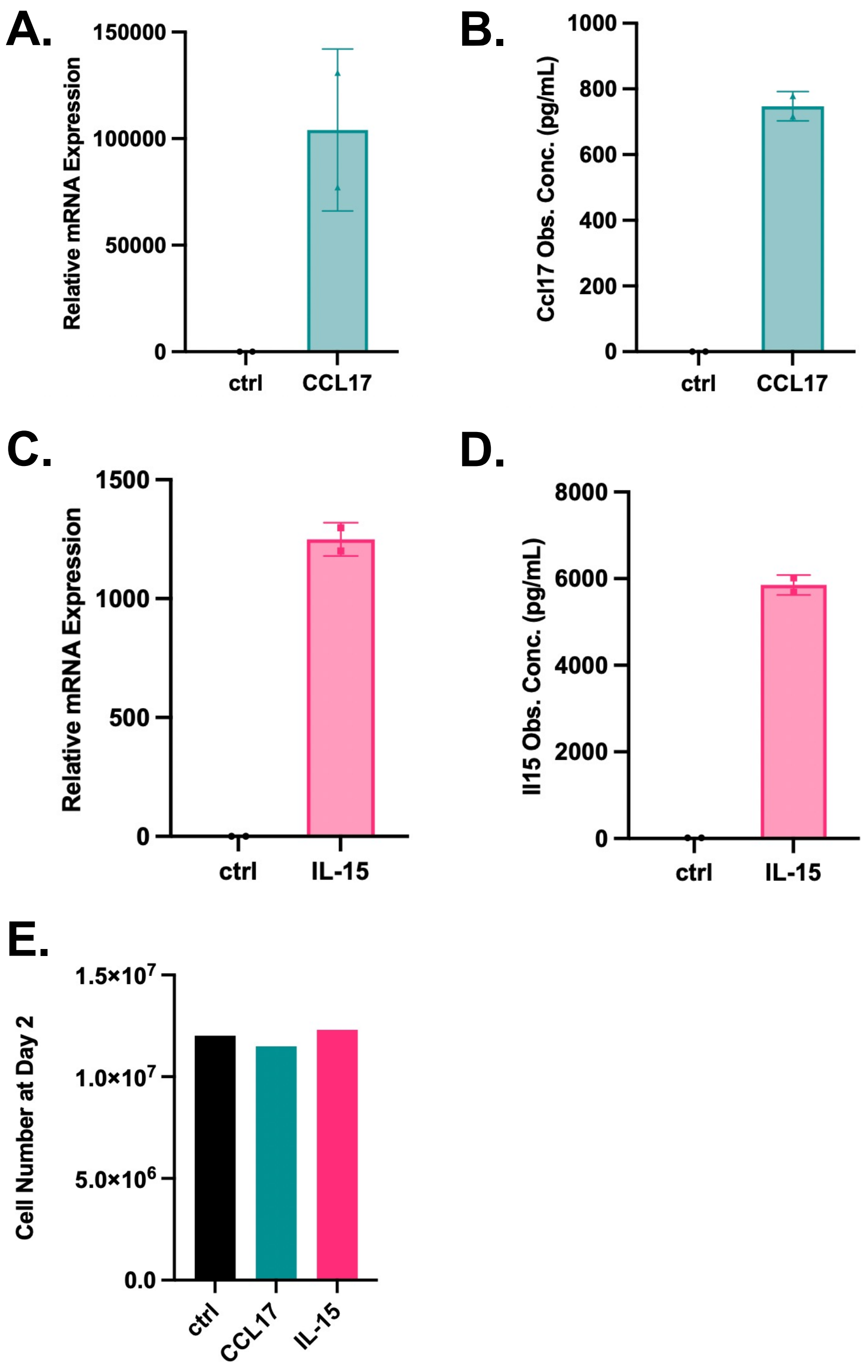


**Fig. S6. Overexpression of mouse *Il15* and *Ccl17* in RIL-175 Ras-driven CCA cell line. (A-B)** qRT-PCR and cytokine array analysis of Ccl17 expression and secretion in RIL-175-Ccl17 compared to control. **(C-D)** qRT-PCR and cytokine array analysis of Il15 expression and secretion in RIL-175-Il15 compared to control. Data are presented as mean ± SD. **(E)** Cell number of control, Ccl17-expressed and Il15-expressed RIL-175 cell lines at day 2. Cells were seeded at $2.2*10^6$ at passage.

### Fig.S8

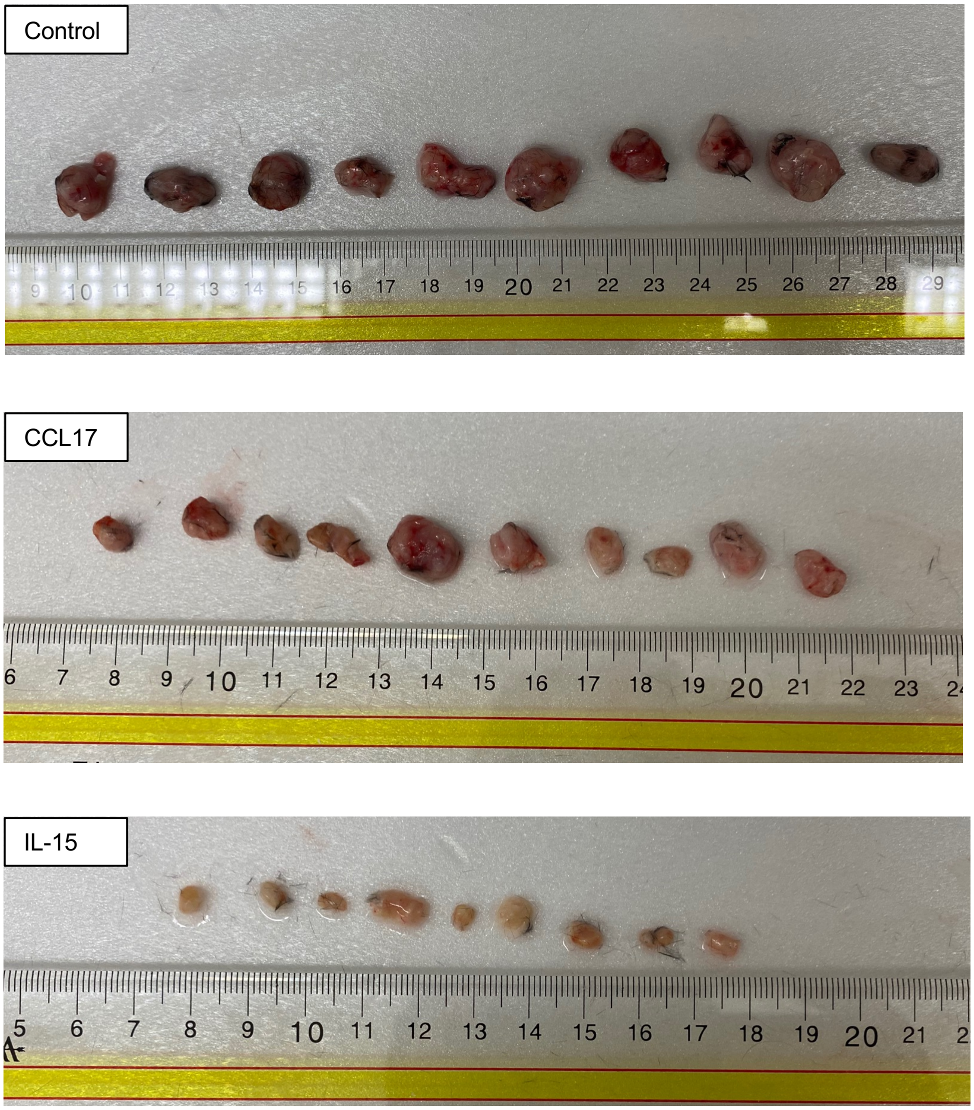


**Fig. S7. Gross tumor appearance of RIL-175 xenograft model at day 13 after implantation.** Tumor appearance of **(A)** RIL-175 control, **(B)** CCL17, and **(C)** IL15 xenograft tumors. Ruler is in cm.

### Fig. S9

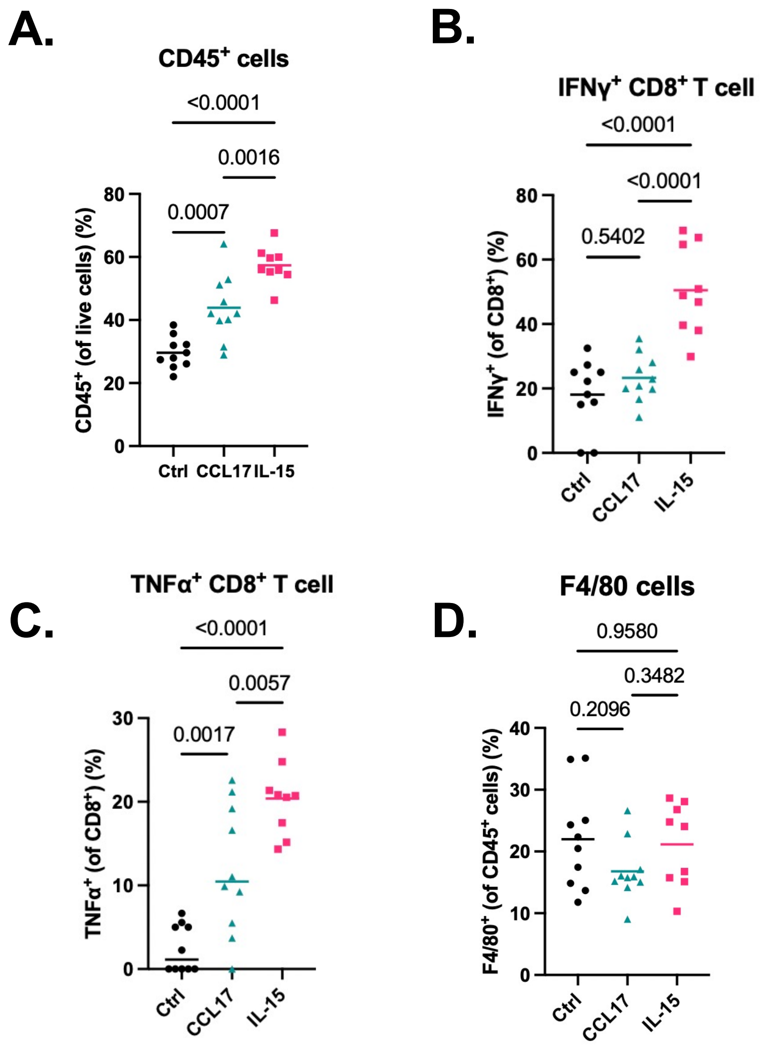


**Fig. S8. Flow cytometry analysis of RIL-175 xenograft tumors at day 13 after implantation. (A)** CD45 positive cells. **(B-C)** IFN$\gamma$ or TNFa positive cells within CD8 positive cell population. **(D)** F4/80 positive cells within CD45 positive cell population. P-values were calculated by one-way ANOVA.
